# A Non-Transcriptional Mitotic Function of POU/Oct Factors Ensures Spindle Stability and Chromosome Segregation

**DOI:** 10.64898/2025.12.19.695428

**Authors:** Priya Gohel, Vasilios Tsarouhas, Laveena Kansara, Suresh Sajwan, Ylva Engström

**Author notes:** Corresponding author/lead contact.

## Abstract

POU/Oct transcription factors are critical regulators of cellular processes, including proliferation, cell fate determination, and cancer. Despite their importance, the specific molecular mechanisms by which they influence cell division remain largely unclear. Here, we show that Nub/Pdm1, a *Drosophila* homolog of human POU2F1/Oct1, is essential for accurate mitotic progression in a non-transcriptional manner. Live imaging and immunostaining in *Drosophila* embryos reveal that its depletion leads to disorganized spindles, aberrant chromosome segregation and delayed mitotic progression. Similarly, reduction of POU2F1/Oct1 in live human cells caused disorganized mitotic spindles and spindle collapse. Nub/Pdm1 is enriched within the mitotic spindles and this recruitment is independent of its sequence-specific DNA binding. Instead, it depends on the integrity of spindle microtubules and is regulated by mitosis-related motor proteins, and kinases. Our findings identify both fly Nub/Pdm1 and human Oct1 as important regulators of mitotic progression, acting to maintain spindle stability and proper elongation. The non-transcriptional mitotic role of Nub/Pdm1 reveals a previously unrecognized mechanism of POU/Oct proteins and provides new insight into their potential oncogenic properties.

**Highlights:**

- Nub/Pdm1 is vital for accurate mitotic progression in a non-transcriptional manner
- Nub/Pdm1 preserves spindle integrity during rapid syncytial nuclear divisions
- Nub/Pdm1 spindle enrichment depends on mitotic factors and intact microtubules
- Nub and human Oct1 ensure proper chromosome segregation

## Introduction

POU/Oct transcription factors are important players in a variety of cellular processes, including proliferation, cell fate determination, immune function, and cancer. It is a versatile group of DNA-binding factors present in all animals (Billin et al., 1991; Bürglin and Affolter, 2016; Herr et al., 1988; McEvilly and Rosenfeld, 1999; Okamoto et al., 1990; Prakash et al., 1992; Rosner et al., 1990; Tang and Engström, 2019; Tantin, 2013; Wegner et al., 1993). Mammalian POU/Oct proteins act as master regulators of pluripotency during embryogenesis, and the pioneer transcription factor POU5F1/Oct4 is pivotal for the maintenance of embryonic stem cells and for pluripotency in mammals. Together with its partners, POU5F1/Oct4 has the capacity to reprogram somatic cells into induced pluripotent stem cells (Okamoto et al., 1990; Soufi et al., 2012). The related POU2F1/Oct1 factor is a ubiquitously expressed transcription factor crucial for embryogenesis and neural stem cell specification, and implicated in a variety of cancers (Vázquez-Arreguín and Tantin, 2016). It serves as the predominant POU/Oct factor in many epithelial tissues, and it has been found to be proto-oncogenic in epithelial malignancies by promoting initiation, progression, and maintenance of epithelial tumors (Jafek et al., 2019; Vázquez-Arreguín et al., 2019).

The *Drosophila* orthologs of the mammalian class II POU factors (POU2F1-3), called Nubbin (Nub)/POU domain protein 1 (Pdm1) and Miti-mere (Miti)/POU domain protein 2 (Pdm2), from here-on referred to as Nub and Pdm2, are known for their crucial involvement in development and lineage specification. Nub and Pdm2 have been implicated in the regulation of both patterning and of precursor cell division in the CNS, and act as key transcription factors in the temporal cascade of lineage specification during embryonic neurogenesis (Doe, 2017; Gabilondo et al., 2016; Tran and Doe, 2008; Tsuji et al., 2008). In adult gut epithelium regeneration, Nub is involved both in stem cell maintenance and in enterocyte lineage differentiation (Tang et al., 2018). Furthermore, Nub, and to some extent also Pdm2, has been shown to be involved in wing and leg specification (Cifuentes and García-Bellido, 1997; Loker and Mann, 2022; Neumann and Cohen, 1998; Ng et al., 1995; Rauskolb and Irvine, 1999).

The *nub* gene is organized in two major transcription units with independent promoters, and these pre-mRNAs produce two distinct protein variants: Nub-PB (long isoform) and Nub-PD (short isoform) (Fig S1A). The common C-terminal region includes the DNA binding POU and Homeo domains. It was recently shown that the two Nub isoforms play antagonistic roles in the regulation of gut immunity, as well as in the control of stemness and differentiation in the regeneration of the gut epithelium (Lindberg et al., 2018; Tang et al., 2018). During *Drosophila* embryogenesis, zygotic *nub* expression is observed as a broad stripe just before cellularization, followed by splitting into two stripes (Affolter et al., 1993; Dick et al., 1991; Lloyd and Sakonju, 1991). Subsequent *nub* expression is dynamic throughout embryogenesis and especially apparent in early neurogenesis when essentially all cells of the neurogenic region express *nub* (Yang et al., 1993). During later stages of embryogenesis, subsets of neuroblasts of the developing CNS express *nub*, as well as cells in the developing anterior and posterior midgut primordia (Affolter et al., 1993). Nub, and to some extent Pdm2, have been implicated in the regulation of a switch between the continued proliferation of stem cells and lineage specification (Bhat and Apsel, 2004; Doe, 2017; Seroka et al., 2020; Tang et al., 2018). However, how Nub acts to maintain proliferation of neuroblasts and intestinal stem cells is not understood.

There is a wealth of studies pointing to the important roles of POU/Oct proteins in regulation of cell cycle progression linked to proliferation and also to cancer (Ben-Batalla et al., 2010; Caelles et al., 1995; Castrillo et al., 1991; Jullien et al., 2015; Shin et al., 2016; Zhao et al., 2014), but few have reached a molecular understanding of the underlying mechanisms. The ubiquitously expressed POU2F1/Oct1 factor is associated with proliferation in cell lines and linked to several cancer types (Jafek et al., 2019; Stepchenko et al., 2022; Vázquez-Arreguín et al., 2019; Vázquez-Arreguín and Tantin, 2016). In addition, POU2F1/Oct1 was found to be regulated by phosphorylation and ubiquitylation in a cell cycle-dependent manner, which affected its interaction with chromatin (Kang et al., 2011).

Here, we intended to gain further mechanistic understanding of the role of POU/Oct proteins in cell proliferation by following cell cycle progression through live imaging in *Drosophila* syncytial embryos, fly and human cell cultures. The well-established pattern of rapid, synchronous nuclear divisions in *Drosophila* syncytial embryos, coupled with the absence of transcription during this period (Edgar and Schubiger, 1986; Harrison et al., 2023; Kwasnieski et al., 2019), makes this model particularly well-suited for exploring putative “non-transcriptional” roles of regulatory proteins. Many transcription and splicing factors end up on mitotic components such as midbody, centrosomes, kinetochores, and spindles during cell division (Kang et al., 2011; Neumann et al., 2010; Pellacani et al., 2018; Somma et al., 2020; Yokoyama et al., 2009), but their roles in mitosis remain elusive.

We found that depletion of Nub-PD isoform, but not Nub-PB, leads to mitotic defects in *Drosophila* S2 cells. In early embryos, loss of Nub-PD causes aberrant anaphase spindles with reduced motility, incorrect chromosome segregation, and mitotic failure, resulting in a significant loss of cortical nuclei and poor embryo survival. These findings suggest that, in syncytial embryos, Nub-PD plays a non-transcriptional role critical for the proper timing and progression of mitosis. Similarly, downregulation of POU2F1/Oct1 in human cells leads to disorganized mitotic spindles and multinucleated cells. Upon mitosis, Nub-PD is enriched around the mitotic spindles, through a regulated mechanism involving components of the microtubule motor proteins Klp61F and Klp3A, the polarity determinant Crumbs, and the kinases Cdk1, Nek2, and NiKi/Nek9. This localization was independent of its sequence-specific DNA binding but depended on spindle microtubule integrity. Together, these findings highlight the critical role of both insect and human POU/Oct factors in mitotic progression, offering new insights into their involvement in normal and malignant cell proliferation.

## Results

### Depletion of Nub-PD perturbs mitosis in *Drosophila* S2 cells

In *Drosophila,* the two Nub protein isoforms, Nub-PB and Nub-PD (Fig. S1A), play antagonistic roles in gut epithelium regeneration, with Nub-PB driving differentiation and Nub-PD maintaining stemness (Tang et al., 2018). How Nub protein isoforms control proliferation/differentiation in opposite directions and how this couples to the cell cycle is not understood. To address this, we first followed the progression of mitosis after isoform-specific RNA interference (RNAi) (Fig. S1B-C). Upon *nub-RB* RNAi, no mitotic phenotypes were observed. In contrast, *nub-RD* RNAi caused striking mitotic defects, with multipolar spindles (34%) and multinucleated cells (32%) compared to control (*luc-RNAi*) (5.5% and 7.2% respectively) (Fig. 1A-B). The presence of multinucleated cells after division was further supported by Lamin staining (Fig. S1D), confirming disrupted nuclear organization after mitosis, suggesting incomplete cytokinesis in these cells. Live imaging of dividing S2 cells expressing histone 2B-GFP (H2B-GFP) and α-tubulin-mCherry, revealed significant delays in metaphase to anaphase transition along with prolonged cytokinesis in *Nub-RD* RNAi cells compared to control cells (Fig. 1C). The delay in mitotic progression was accompanied by defects in spindle integrity in *nub-RD* RNAi cells. Following the cell-cycle phase distribution in Fly-FUCCI (Fluorescent Ubiquitination-based Cell Cycle Indicator) S2 cells (Zielke et al., 2014) showed that the proportion of cells in S phase was unaffected by the knockdown of either isoform (Fig. S1E). However, *nub-RD* knockdown significantly increased the number of G2/M phase cells and reduced G1 phase cells, indicating a block in mitotic entry or progression. In contrast, *nub-RB* knockdown slightly reduced G2/M phase cells and increased G1 phase cells (Fig. S1E), suggesting distinct roles for the two Nub isoforms during the cell cycle. Collectively, these results indicate an important role of Nub protein in cell division, with the entry and/or progression through mitosis specifically requiring the Nub-PD isoform.

**Figure 1.**
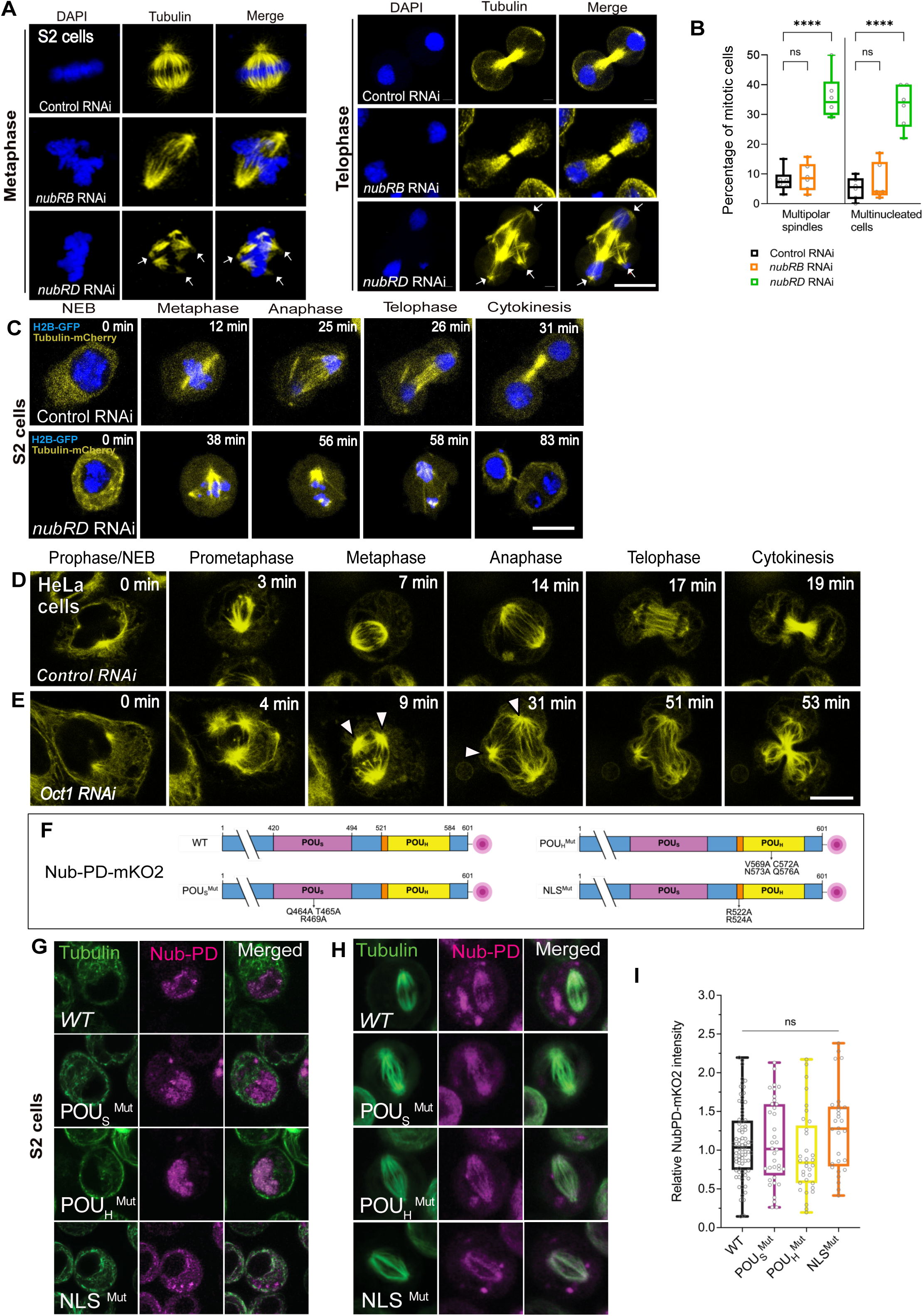
RNAi-mediated depletion of Nub-PD causes mitotic spindle defects in *Drosophila* S2 cells, independently of its nuclear function. (A) Confocal images of mitotic S2 cells in metaphase and telophase, treated with dsRNA as indicated (RNAi) and processed for immunofluorescence with β-tubulin (yellow). DNA was counterstained with DAPI (blue). White arrows indicate abnormal spindle morphology. (B) Box and whisker graph showing mitotic defects (multipolar spindles and multiple nuclei) in control, *nub-RB RNAi,* and *nub-RD* RNAi-treated cells. Statistical significance was calculated by one-way ANOVA followed by Tukey’s multiple comparison test. *ns* = not significant; * *p < 0.05*, *\*\*\*\* p < 0.0001*. (*N* = 6 biological replicates and *n* = 620 mitotic cells). (C) Representative images from live imaging of S2 cells stably expressing H2B-GFP (blue) and α-tubulin-mCherry (yellow). Mitotic progression of control and *nub-RD* RNAi-treated cells, relative time (min) as indicated. (D-E) Representative images of live HeLa-cells treated with control (D) or POU2F1/Oct1*-*siRNA (E) and incubated with ViaFluor^®^ 488 dye to visualize the spindle microtubules (yellow). The relative time (min) during mitotic progression is indicated. (F) Graphical representation of Nub-PD-mKO2 protein domains of wild type (WT) and mutated Nub-PD variants, tagged in the C-terminus with monomeric mKO2 (red circle). WT: full length Nub-PD-mKO2 protein; POU_S_^Mut^: mutated POU-specific (POU_S_) domain at Q464A, T465A and R469A; POU_H_^Mut^: mutated POU homeo domain (POU_H_) at V569A, C572A, N573A and Q576A; NLS^Mut^: mutated nuclear localization signal (NLS) at R522A and R524. (G-H) Confocal images of live S2 cells during interphase (G) and metaphase (H) transfected with the NubPD-mKO2 plasmids. Cells were incubated with ViaFluor^®^ 488 dye to visualize the spindle microtubules (MTs) (green). The mKO2 signals (magenta) are overlapping with MTs (green) for the wt and all mutated variants. (I) Graph showing the relative spindle intensity of mKO2 in wt and mutated variants of NubPD-mKO2. The box plots show the median (horizontal line), and the data range from 25^th^ to 75^th^ percentiles. The bars indicate maximum and minimum values. *ns* denotes no statistical significance, as calculated by unpaired two tailed *t*-tests.

### POU2F1/Oct1 is required for mitotic progression in human cells

To examine whether a mitotic function is evolutionarily conserved among POU/Oct factors, we investigated a similar role of the human homologue, POU2F1/Oct1, in mitotic progression. Using siRNA-mediated knockdown of POU2F1/Oct1 in HeLa cells, combined with live-cell imaging, revealed that POU2F1/Oct1 depletion causes prolonged mitosis, disorganized spindle structures, and cytokinesis failure (Fig. 1D-E). These results were consistent with quantitative experiments in fixed HeLa and HEK293 cells, with a 3.5-fold increase multinucleated cells (28.2%) compared to controls (6.7%, p < 0.01) (Fig. S1F–G), and a 3.8-fold increase in HEK293 cells (28.6% vs. 7.8%, p < 0.01) (Fig. S1H–I). These findings are similar to those seen upon Nub-PD downregulation in *Drosophila* S2 cells (Fig. 1A-C), suggesting an analogous and possibly evolutionarily conserved role of these POU/Oct proteins in mitotic timing and progression.

### Sequence-specific DNA binding is not required for mitotic spindle localization of Nub-PD

To further investigate the mitotic role of Nub, we generated Nub-PD constructs tagged with monomeric Kusabira-Orange 2 (mKO2) and Red Fluorescent Protein (RFP), and analyzed their intracellular localization by live imaging in *Drosophila* S2 cells. Nub-PD-mKO2 and Nub-PD-RFP localized to the nucleus during interphase (Fig. 1G, Fig. S1K–L). During mitosis, however, Nub-PD-mKO2/RFP localized around the mitotic spindle (Fig. 1H–I, Fig. S1K), overlapping with tubulin staining, further supporting a mitotic role. Next, we explored if the localization around the mitotic spindle involved amino acids required for transcriptional regulation or not. Nuclear localization and DNA binding properties of the POU and Homeo domains of POU/Oct factors have been extensively studied (Phillips and Luisi, 2000; Sturm and Herr, 1988).

Structure-function analyses have pinpointed the amino acids that are crucial for sequence-specific DNA binding and transcriptional regulation (Assa-Munt et al., 1993; Dekker et al., 1993; Klemm et al., 1994; Sturm et al., 1988). The DNA binding POU specific (POU_S_) and POU Homeo (POU_H_) domains of *Drosophila* Nub and human POU2F1/Oct1 show 95% and 83% identity in the POU_S_ and POU_H_ domains respectively. Building on this knowledge and high amino acid conservation we expressed tagged Nub-PD protein variants with mutations in the POU_S_ and POU_H_ domains that are identical in the two proteins, and shown to be crucial for sequence-specific DNA binding of POU2F1/Oct1(Klemm et al., 1994). We mutated the POU_S_ domain at Q464A, T465A and R469A; the POU_H_ at V569A, C572A, N573A and Q576A; and the nuclear localization signal (NLS) at R522A and R524A (Fig. 1F). As expected, mutations in the NLS blocked nuclear localization of NubPD-mKO2 in interphase cells (Fig. 1G and Fig. S1M), while mutations in the POU and Homeo domains did not (Fig. 1G). In metaphase cells, both the NLS mutant and the Nub-PD Homeo and POU domain mutants localized to the mitotic spindle (Fig. 1H-I), indicating that sequence-specific DNA binding is not required for this localization. Thus, sequences in Nub-PD required for nuclear targeting and transcriptional regulation are not critical for the localization to the mitotic spindle, and therefore unlikely to be involved in the mitotic functions of this protein.

### Nub-PD is essential for mitosis and nuclear divisions during early ***Drosophila* embryogenesis**

To examine the mitotic functions of the Nub-PD isoform *in vivo* during development we utilized the syncytial blastoderm stage of *Drosophila* embryos, which entirely relies on maternally provided transcripts due to the absence of transcription (Fig. S2A). Early *Drosophila* embryogenesis is characterized by rapid and synchronous nuclear divisions switching between S and M phases, creating a syncytial blastoderm (Fig. S2A) (Harrison et al., 2023). Previous studies have shown *nub* mRNA expression in syncytial embryos (Lécuyer et al., 2007; Wilk et al., 2016). Consistently, robust expression of *nub-RD*, but not of *nub-RB*, was observed in 0-2 h embryos (Fig. S2B). Unfertilized oocytes dissected from female ovaries revealed no Nub protein immunostaining, indicating that *nub* mRNA is not translated during oogenesis (Fig. S2C). However, Nub immunostaining was evident throughout newly laid embryos at nuclear division cycle 1 (NC1) (Fig. S2D), and in nuclei at NC7 and NC12 before zygotic genome activation (ZGA) (Fig. S2F), demonstrating efficient translation of *nub* mRNA during the syncytial stages. Furthermore, embryos laid by females mated with “spermless” *tudor (tud)* males also expressed Nub protein (Fig. S2E), reinforcing that *nub is* expressed from the female germline and prior to ZGA. It has been previously reported that a large fraction of maternally provided transcripts are translated during ovulation in a process called egg activation (Harrison et al., 2023; Krauchunas and Wolfner, 2013). Our results demonstrate that *nub* is one of those genes that is maternally transcribed and subsequently translated during egg activation. Importantly, we observed strong Nub immunostaining around the mitotic spindle of the first nuclear division (Fig. S2D-F, arrowheads) suggesting that Nub-PD has a function in the very first and all subsequent nuclear divisions of the syncytial embryo.

Next, we investigated the mitotic progression during early embryogenesis in embryos with reduced Nub-PD levels using a recessive hypomorphic mutation, *nub^1^*, with strongly diminished *nub-RD* expression due to a transposon located just upstream of the *nub-RD* transcription start site (Fig. S2G-I’’) (Ng et al., 1995). Despite a previous report showing diminished expression of the *pdm2* paralog in *nub^1^* mutant wing discs (Loker and Mann, 2022) we found no effect on Pdm2 protein expression in *nub^1^* syncytial embryos (Fig. S2J). Homozygous *nub^1^* females are fertile, but the embryo hatch rate was only 60% (Fig. S2K), demonstrating compromised development. This enabled us, however, to analyze mitosis in early syncytial embryos with reduced Nub-PD levels. By following nuclear divisions in *nub^1^*embryos, we noticed disrupted nuclear distribution in the cortex with areas devoid of nuclei, especially at NC10-11 (Fig. 2A-B), and reduced nuclear occupancy at NC10-13 compared to control embryos (Fig. 2C). Further, in both *nub^1^* mutants and *MTD>nub RNAi* embryos, we observed normal chromosome alignment at metaphase (Fig. 2D-E) but obvious chromosome mis-segregation during anaphase, including abnormal anaphase chromosome geometry. These defects persisted into telophase as unresolved chromosome bridges (Fig. 2F-I), leading to nuclear fallout (NUF), a well-described endpoint for defective nuclei (Rothwell et al., 1998; Takada et al., 2003). Although partial recovery in nuclear occupancy was observed in some *nub^1^* embryos at NC12-13 (Fig. 2C), we conclude that the pronounced division failure is likely a major cause of the poor embryo survival and low hatch rate of *nub^1^* mutants (Fig. S2K).

**Figure 2.**
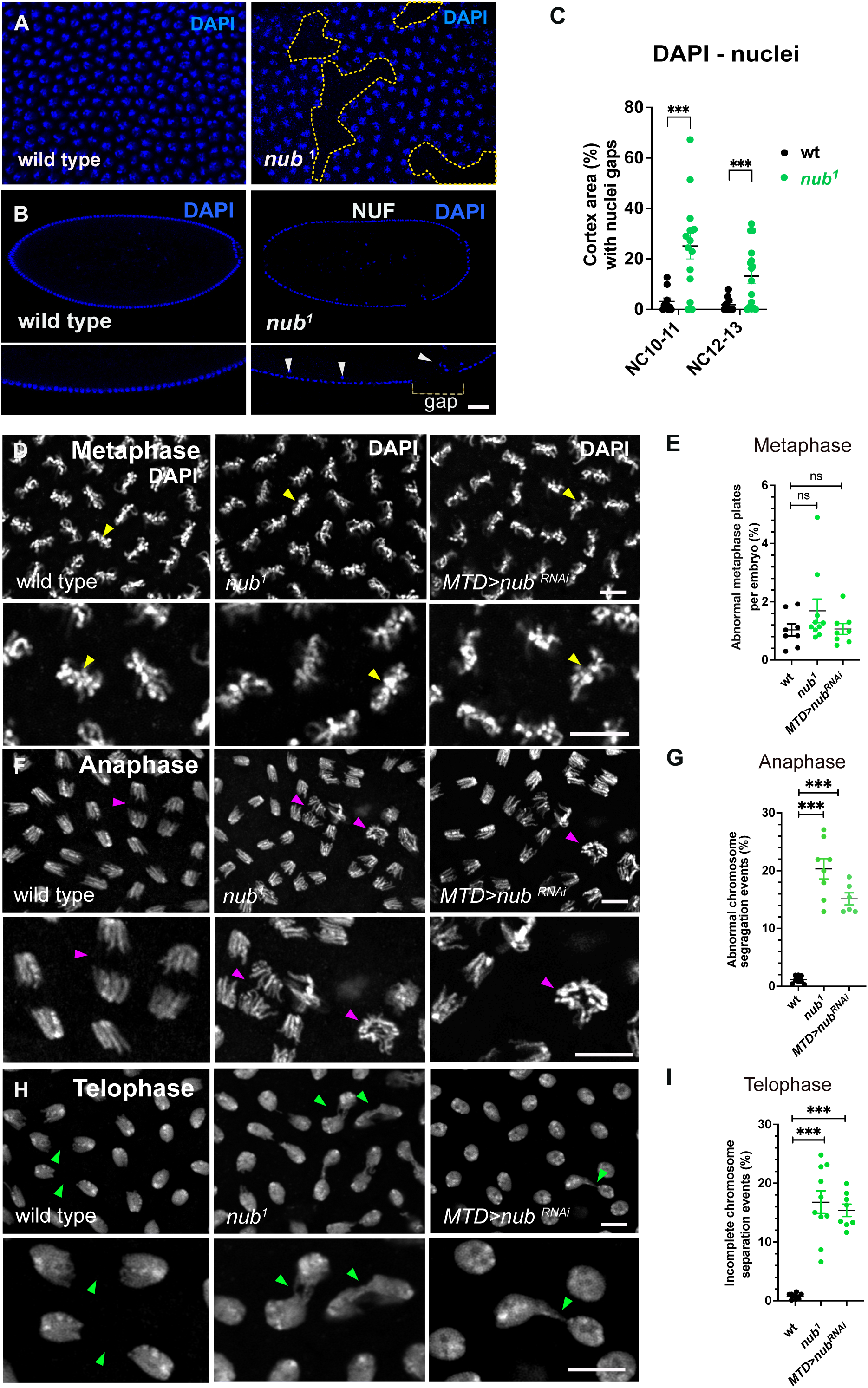
Nub is required for efficient nuclear division and faithful chromosome segregation in *Drosophila* syncytial embryos. (A) Confocal images displaying the cortex of wild-type and *nub^1^* embryos stained with the nuclear probe DAPI. Nuclei-free cortical areas (dashed lines). (B) Confocal middle sections of wild-type and *nub^1^* embryos stained for DAPI. Images below are zoomed-in areas of the cortex, shown in the upper images. Arrowheads depict nuclei falling into the inner yolk region. (C) Scatter plot showing the percentage of the cortical area with nuclei gaps in wild-type (*n* = 18) and *nub^1^* (*n* = 27) mutant embryos during indicated nuclear cycles (NC). *** indicates *p < 0.001,* calculated by unpaired two-tailed t-tests. (D-H) Representative confocal images of *Drosophila* pre-blastoderm embryos at metaphase (D), anaphase (F), and telophase (H) stained with DAPI to visualize chromosomes. Wild-type embryos show properly aligned metaphase chromosomes (yellow arrowheads), normal segregation during anaphase (purple arrowheads), and evenly sized daughter nuclei in telophase (green arrowheads). In contrast, *nub¹* mutant and *MTD>nub RNAi* embryos display chromosome misalignment, lagging chromosomes, and unequal chromosome distribution, indicating disrupted mitotic progression. (E-I) Quantification of abnormal chromosome configurations in metaphase (E), anaphase (G), and telophase (I). The percentage of nuclei showing misaligned chromosomes (metaphase), abnormal chromosome segregation (anaphase), or incomplete/unequal segregation (telophase) was significantly increased in nub¹ and MTD>nub RNAi embryos relative to wild-type. Each data point represents one embryo; mean ± SE shown. Statistical significance was assessed using *t*-test; ns, not significant; ***P < 0.0001. (Scale bars: 10 μm).

To further explore the role of Nub-PD in nuclear divisions we performed time-lapse microscopy during the last four syncytial mitotic cycles (NC 10-13) in wild-type and *nub^1^* embryos expressing the RFP-tagged histone H2A variant His2Av (His2A-RFP)(Pandey et al., 2005). First, we investigated if the defective nuclear divisions were the result of aberrant DNA synthesis (S-phase) or mitosis (M-phase). During syncytial nuclear divisions, nuclei alternate between S/M phases, with no gap phases (G-phases) (Glover, 1991; Harrison et al., 2023) (Fig. S2A). The duration of the S phase was indistinguishable between *nub^1^* and control embryos (Fig. 3A, B), but the M phase was significantly longer in *nub^1^* mutant embryos in NC 11-13 (Fig. 3A, C). Additionally, time-lapse imaging revealed five mitotic defects in *nub^1^* embryos: clustered/aggregated nuclei, asymmetrically spread nuclei, asynchronous divisions, arrested divisions, and cortical gaps with NUF (Fig. 3D-F, Fig. S3A-B, S3E and Videos S1-3). Approximately 55% of *nub^1^* embryos displayed clustered/aggregated nuclei, and 45% exhibited large NUF patches, consistent with the phenotypic analysis based on immuno-staining of fixed embryos. Only 12-16% of wild-type controls showed such nuclear-division phenotypes. Moreover, *nub^1^* embryos showed significant asynchrony and failed to maintain spatial nuclear arrangements, leading to asymmetric spreading of nuclei in 25% of embryos (Fig. S3A-C). Small subsets of *nub^1^* embryos (15 %), compared to wild type (3%), were arrested and failed to progress with subsequent nuclear divisions (Fig. S3B).

**Figure 3.**
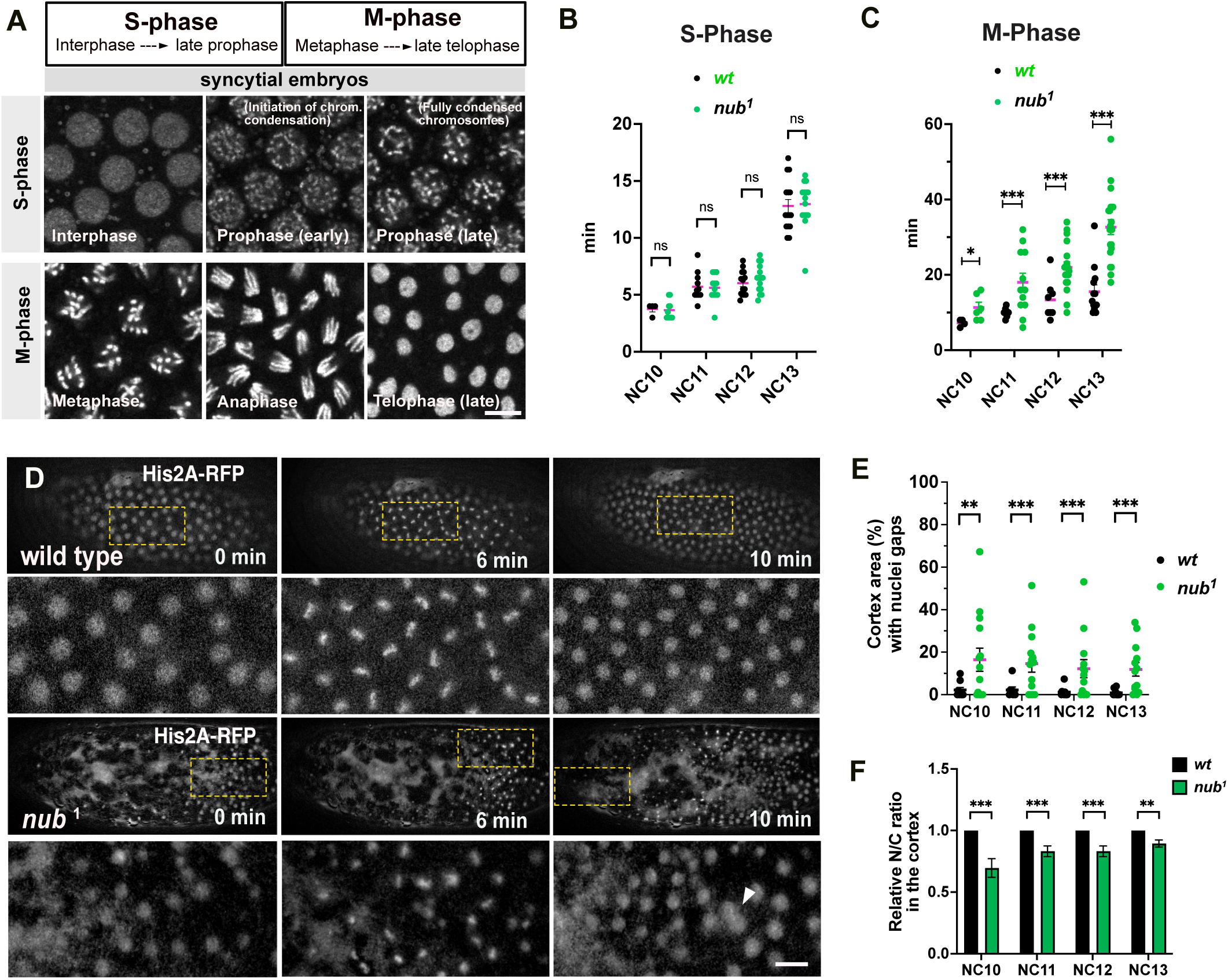
Loss of Nub disrupts the timing and organization of nuclear divisions during early syncytial development. (A) Reference panel showing DAPI-stained nuclei from *Drosophila* syncytial embryos, used to define S-phase (interphase to late prophase) and M-phase (metaphase to late telophase) during nuclear cycles (NC10–13). The phases were classified according to (Fasulo et al.). Scale bar: 10 µm. (B-C) Scatter plots showing the time in minutes (min) spent in S phase (B) and M phase (C) in wild type *(n* = 48) and *nub^1^ (n* = 85) embryos during NC10-13. *ns* denotes no statistical significance, * *p > 0.05*; *** *p < 0.0001,* as calculated by unpaired two tailed *t*-tests. (D) Fluorescent images from time-lapse recordings showing live wild type and *nub^1^* mutant embryos at NC10-11. Embryos are expressing the His2A-RFP reporter. Images below depict zoomed-in areas of the embryo cortex indicated by the rectangular frames in the upper images. (E-F) Plots showing the percentage of cortical area with nuclei gaps (E), or the relative nuclear to cytoplasmic ratio (N/C ratio) (F), per embryo in wild-type (*n* = 47) and *nub^1^* (*n* = 57) mutant embryos during different nuclear cycles (NC10-13). Values were normalized to wild type. **, *** denote statistical significance *p* < 0.01, *p* < 0.001, respectively, as calculated by unpaired two tailed *t*-tests.

### Loss of Nub-PD affects spindle organization and anaphase elongation

To understand the primary cause of the phenotypes of dividing nuclei upon Nub-PD protein loss, we examined the mitotic spindle organization by co-immunostaining of β - and γ-tubulin in wild type and *nub^1^* embryos. As shown in Fig. 4A-B and Videos S5-6, *nub^1^* embryos displayed disorganized spindles with detached microtubule bundles, particularly during the NC11-13 of syncytial development. We also frequently observed free centrosomes at the cortex, and spindles disconnected from centrosomes below the cortex, which then fell into the interior of the embryo for spindle MT de-polymerization (Fig. 4B-C).

**Figure 4.**
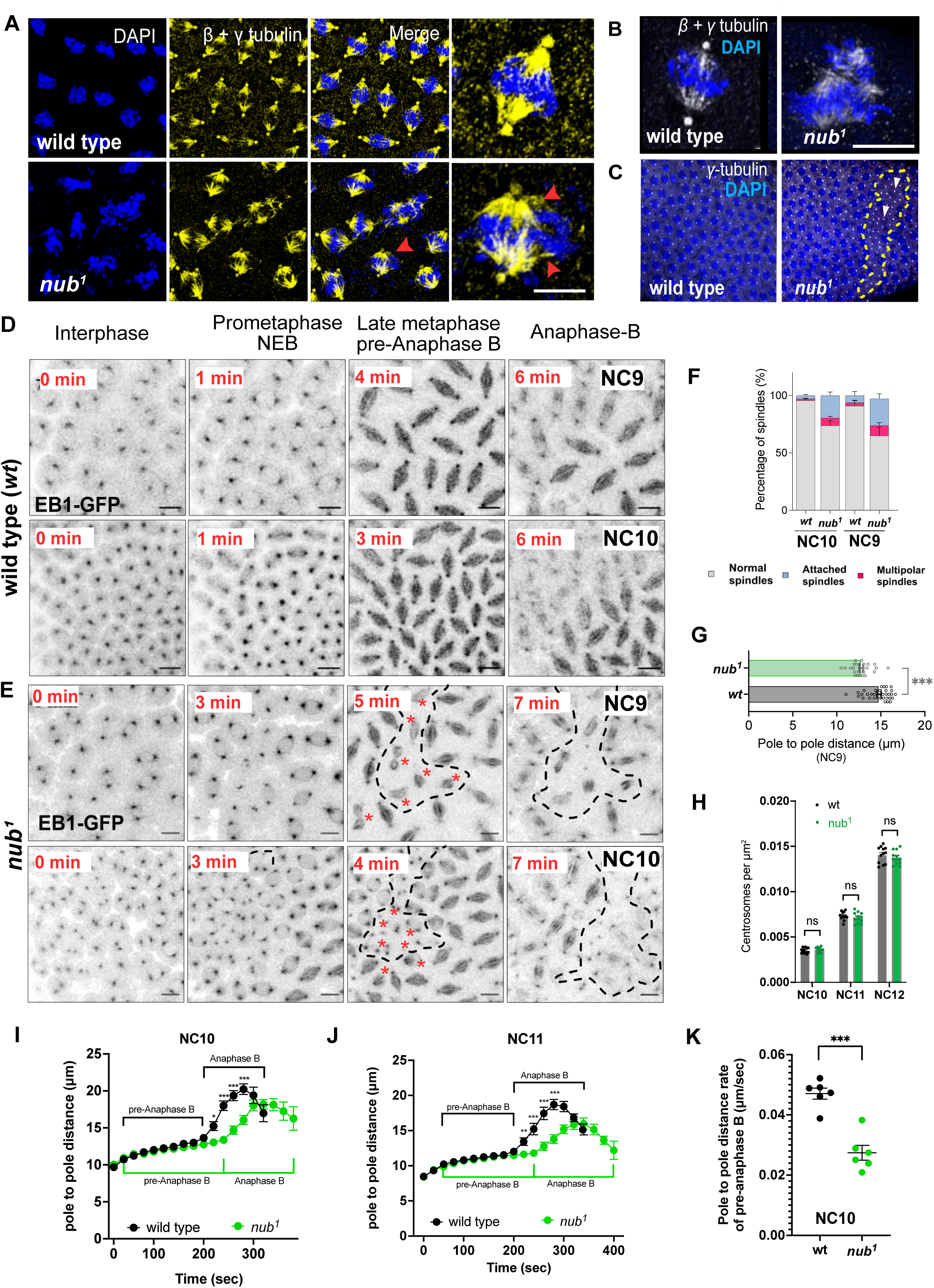
Nub-PD is necessary for proper spindle organization and elongation in syncytial-blastoderm stage embryos. (A) Representative images showing *w^1118^* and *nub^1^* syncytial embryos stained for β-tubulin, γ-tubulin (yellow) and DNA with DAPI (blue). Disorganized *nub^1^*spindles with detached microtubular bundles (red arrows) are shown (lower enlarged image). (B) Confocal images displaying single spindles of wild-type and *nub^1^* mutant embryos stained for β*-*tubulin (spindles, white) and γ-tubulin (centrosomes, white) and DNA with DAPI (blue). (C) Confocal images of the cortex of wild-type and *nub^1^* mutant embryos stained for γ*-*tubulin (white) and DNA with DAPI (blue). Cortical areas with centrosomes and no nuclei are indicated (dash lines). Arrowheads denote free centrosomes. (D-E) Representative time-lapse images of wild type (D) and *nub^1^* mutant (E) embryos expressing EB1-GFP, in NC9 and NC10. The corresponding time (min) from interphase to anaphase is indicated. Dotted areas, along with asterisks, indicate groups of disorganized or attached spindles and free centrosomes. (F) The percentage of normal, multipolar and attached spindles is shown for *wt* and *nub^1^* mutant embryos at NC9 and NC10. *n* = 30-50 mitotic spindles, *N* = 5 embryos. Statistical significance was assessed by two-way ANOVA (multiple comparison analysis). (G) Quantification of centrosome density per μm^2^ at nuclear cycles NC10–12 showing no significant difference between wild-type and *nub¹* embryos. Mean ± SE; ns, not significant. Statistical significance was calculated by *t*-test. (H) Spindle length (pole-to-pole distance) in *nub¹* and wild type embryos at NC9. Mean ± SD; ***P < 0.001. (I-J) Pole-to-pole distance (μ*m*) as a fraction of time (sec) in wt and *nub^1^* dividing spindles expressing EB1-GFP during metaphase, pre-anaphase B and anaphase B at NC10 (I) and NC11 (J). Each time point represents the average data from multiple spindles (n) and embryos (N). *n* = 128-137, *N* = 9-10 embryos per genotype. The statistical significances (3H and 3I) were obtained from a logistic regression model using the *ChiSq* test. *, *p < 0.05*, ***, *p < 0.0001*. (K) Graph showing the average rates of spindle pole-to-pole distance (μ*m*/sec) in *wt* (*N* = 6 embryos, *n* = 78 spindles) and *nub^1^* (*N* = 6 embryos, *n* = 67 spindles) mutant embryos expressing EB1-GFP. ***, *p < 0.0001.* Statistical significance was calculated by unpaired two tailed *t*-test. Scale bars, 10 μm.

Live-imaging of embryos expressing the microtubule plus-end binding protein 1 (EB1) tagged with GFP (EB1-GFP)(Piehl et al., 2004) showed normal spindle dynamics in control embryos (Fig. 4D), as previously reported (Karr and Alberts, 1986; Kellogg et al., 1988; Sullivan and Theurkauf, 1995). In contrast, *nub^1^* embryos produced mitotic spindles during metaphase but failed to maintain their proper size and shape (Fig. 4D-F). Pre-anaphase B spindle length (pole-to-pole distance) was shorter in *nub^1^* embryos compared to control embryos (Fig. 4G) at NC9. Spindles progressively collapsed and detached from centrosomes in metaphase and anaphase, leading to accumulation of free centrosomes at the cortex (Fig. 4D-E), consistent with the γ-tubulin staining (Fig. 4B-C). The centrosome density (centrosomes per µm^2^) was unchanged in *nub^1^* mutants compared with wild type, arguing against defects in centriole duplication or centrosome amplification (Fig. 4H).

Tracking individual spindles in EB1-GFP embryos revealed significant delays in the metaphase-to-anaphase transition in *nub^1^* embryos at both NC10 and NC11 (Fig. 4I-J), consistent with the defects observed at NC9. Wild-type spindles showed dynamic changes in pre-anaphase B with an average elongation rate of 0.047 μm/sec. In contrast, *nub^1^* mutant spindles elongated more slowly, averaging 0.027 μm/sec (Fig. 4K). These phenotypes suggest a role of Nub-PD in anaphase spindle elongation. Taken together, our data indicate that Nub-PD plays a pivotal role in spindle organization and elongation to sustain rapid syncytial nuclear divisions.

We then aimed to restore the mitotic function in *nub^1^* genetic background by introducing a functional *nub* gene copy via a *nub* locus duplication on the 3rd chromosome (CH321-40A01). The duplication line showed increased Nub immunostaining compared to wild type in zygotic embryos (Fig. S4A). The *nub* locus duplication provided a significant rescue of the NUF phenotype (Fig. S4B-C), and of the mitotic spindle organization in *nub^1^* mutants (Fig. S4D-E). These results confirm that the mitotic failures were specifically due to the loss of *nub*, and highlight the essential contribution of Nub-PD for spindle stability, and consequently for proper chromosome segregation during mitosis.

### A critical and non-transcriptional role of Nub-PD in early embryos

To further validate the essential role of Nub-PD protein in the early embryo we specifically depleted Nub-PD protein without affecting the maternally loaded *nub-RD* mRNA, using targeted degradation of GFP-fusion proteins (deGradFP)(Caussinus et al., 2011) (Fig. 5A). We first created Nub-PD and Nub-PB tagged proteins endogenously with the superfolder Green Fluorescent Protein (sfGFP), and then generated two independent transgenic fly lines (*sfGFPnub-PB* and *sfGFPnub-PD*) by CRISPR/Cas9-mediated homology-directed repair (HDR) (Fig. 5B). Both sfGFP-tagged transgenic lines were homozygous viable and fertile. Further, the reduced nuclear-to-cytoplasmic (N/C) ratio at NC11–12 in *nub¹* mutants was returned back to wild type levels in heterozygous *sfGFP-nub-PD/ nub¹* embryos generated by introducing the *sfGFP-nubPD*-allele into the *nub¹* background (Fig. S5A). The adult life span of *sfGFP-nubPD* and *sfGFP-nub-PB* was near wild-type levels (Fig. S5B), demonstrating that the tagged protein forms are functionally competent *in vivo*. Consistent with the mRNA measurements (Fig. S2B), early *sfGFP-nubPD* embryos, but not *sfGFP-nubPB* embryos showed a GFP signal (Fig. 5C-D).

**Figure 5.**
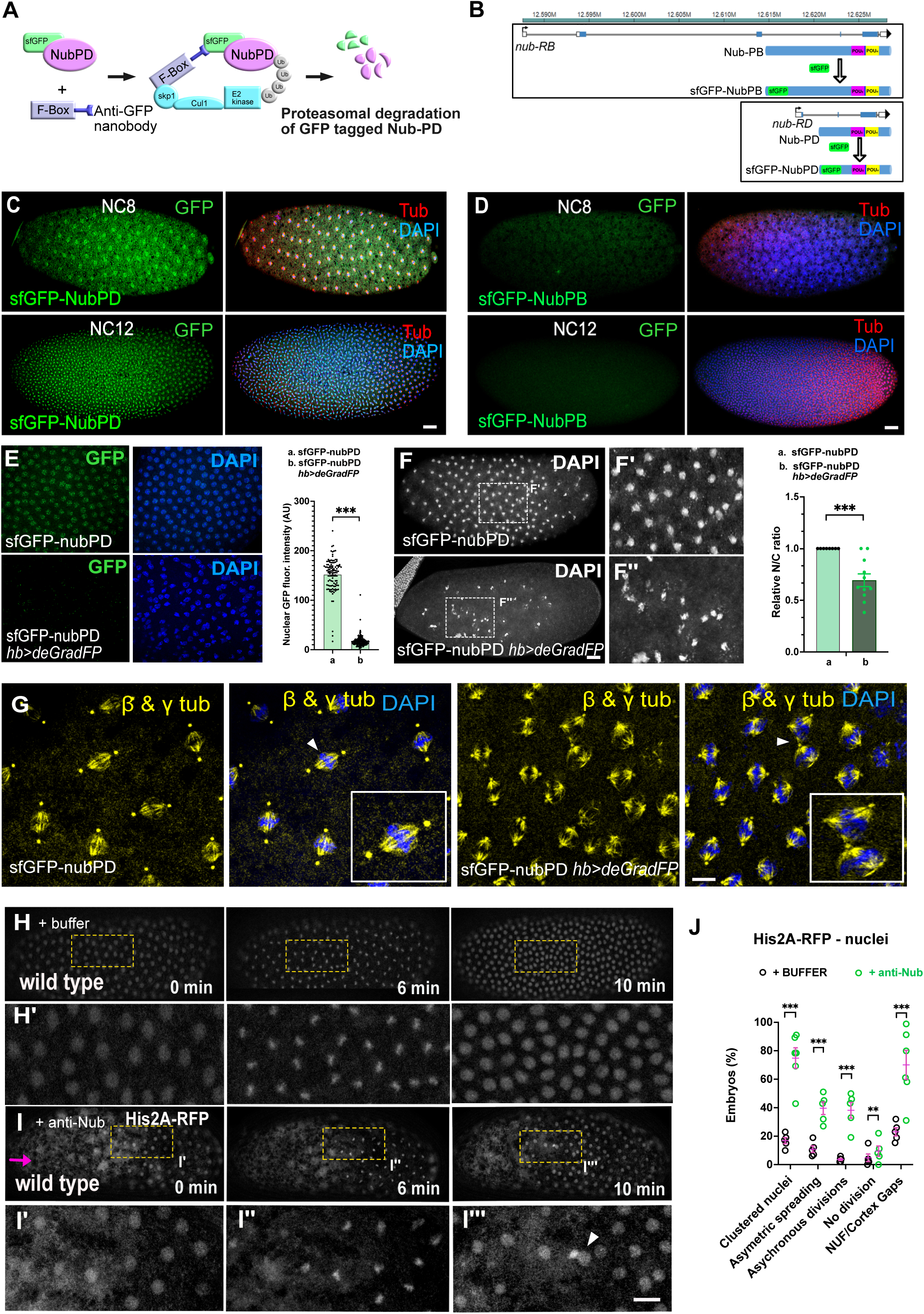
Impairment of Nub-PD in syncytial embryos demonstrates a non-transcriptional role in mitosis. (A) Schematic representation of the deGradFP (utilizing ubiquitin-proteasome pathway) for degradation of GFP-tagged Nub-PD protein in embryos. (B) Schematic representation of sfGFP-tagged Nub protein isoforms, sfGFP-NubPB and sfGFP-NubPD. Independent fly lines were generated by CRISPR-Cas9-mediated genome editing, tagging endogenous Nub-PB and Nub-PD proteins with super folder Green Fluorescent Protein (sfGFP). (C) Fluorescent images of *sfGFP-NubPD* (green) in embryos at NC8 and NC12. β-tubulin (red) and DAPI (blue). (D) Fluorescent images of *sfGFP-NubPB* (green) in blastoderm staged embryos at NC8 and NC12. β-tubulin (red) and DAPI (blue). (E) Representative confocal images of control *sfGFP-Nub-PD* embryos (top) and embryos expressing *hb>deGradFP* (bottom), stained with DAPI and antibodies against GFP. Quantification (right) shows a strong reduction in nuclear GFP intensity (Arbitrary Units, A.U.) upon *deGradFP* expression. *p* < 0.001, calculated by unpaired two tailed *t*-test. (F) Fluorescent images of control (*sfGFP-NubPD)* and *sfGFP-NubPD*; *hb>deGradFP* embryos during NC8, stained for DNA by DAPI (white). Quantification (right) reveals a significant reduction in nuclear to cytoplasmic (NC) ratio upon *sfGFP-NubPD* depletion. *p* < 0.0001, calculated by unpaired two tailed *t*-test. (H) Fluorescent images from time-lapse recordings showing live wild type buffer-injected embryo at NC10-11. Embryos are expressing the His2A-RFP reporter. Images below depict zoomed-in areas of the embryo cortex indicated by the rectangular frames in the upper images. (I) Fluorescent images from time-lapse recordings showing live wild-type embryos injected with anti-Nub antibody. Embryos are expressing the His2A-RFP reporter. Images below indicate zoomed-in areas of the embryo cortex indicated by the rectangular frames in the upper images. The injection site is indicated by an arrow (magenta). Arrowhead indicates abnormal nuclei division. (G) Confocal images showing the organization of the mitotic spindles in control (*sfGFP-NubPD*) and *sfGFP-NubPD*; *hb>deGradFP* embryos. Tubulin (β*-* tubulin *+* γ-tubulin, yellow) and DAPI (blue).

Next, the *sfGFPnub-PD* line was crossed with the previously described *hunchback*-deGradFP (*hb*-deGradFP) and *nanos*>deGradFP (*nos>*deGradFP) transgenic lines(Gaskill et al., 2021; Vazquez-Pianzola et al., 2022). The *hb*-minimal maternal promoter and the *nos* promoter drive the expression of *deGradFP* mRNA during oogenesis, which subsequently is translated into active deGradFP protein upon egg activation/ovulation, leading to degradation of newly translated sfGFP-tagged proteins in early embryogenesis. The sfGFP-NubPD fluorescence was strongly reduced by expression of the *deGradFP* in early syncytial embryos, confirming prominent degradation of GFP-tagged Nub-PD protein (Fig. 5E). These Nub-PD-depleted embryos exhibited asynchronous nuclear divisions and chromosome segregation defects (Fig. 5F-F’’ and Fig. S5C). Additionally, degradation of sfGFP-NubPD caused severe spindle organization phenotypes, characterized by detached microtubule bundles similar to those observed in *nub^1^* mutants (Fig. 5G). This confirms that loss of Nub-PD in transcriptionally silent syncytial embryos, and not during oogenesis, when transcription is functionally robust, causes the aberrant mitotic phenotypes.

To further test the non-transcriptional function of Nub in nuclear divisions, we acutely inhibited Nub protein in wild-type embryos by microinjecting an anti-Nub antibody during early syncytial nuclear cycles. Live imaging of wild-type embryos expressing His2A-RFP, micro-injected at NC9 with an antibody against Nub protein, recapitulated the effects observed in *nub^1^* embryos (Fig. 5H-I’’’ and S5D). Antibody injection led to a 75% incidence of clustered/aggregated nuclei, compared to 15% in controls, causing chromosome segregation phenotypes (Fig. 5J and Video S4). These phenotypes were particularly strong near to the injection site, marked by labelled dextran (3 kDa). As expected, the nuclei from these highly irregular divisions subsequently fell into the interior of the embryo (NUF phenotype), creating large cortical nuclei gaps (Fig. 5I-I’’’, and Fig. S5D). Taken together, our data demonstrate that Nub-PD is required for an essential non-transcriptional function in mitotic progression and nuclear division *in vivo*.

In *nub^1^* embryos, mitotic defects were prominent at NC10-13 but less evident at earlier NCs (Fig. S5F), despite that Nub localized to the mitotic spindle during the first nuclear division (Fig. S2F). We hypothesized that the less pronounced phenotype at early NCs reflects residual Nub-PD protein in the hypomorphic *nub^1^* allele (Fig. S2I-I’’). To investigate this, we combined the *nub^1^* mutant with *nub* RNAi knockdown using the maternal driver matα4-Gal4-VP16. This led to more severe mitotic defects, including chromosome segregation errors (Fig. S5E-G) and spindle organization defects (Fig. S5H-I) as early as NC5, supporting the hypomorphic nature of the *nub^1^* allele. Given that the earliest signs of zygotic gene transcription have been reported at NC8 (Edgar and Schubiger, 1986; Erickson and Cline, 1993; Kwasnieski et al., 2019; Pritchard and Schubiger, 1996), the mitotic defects recorded here at NC5 further argues for a non-transcriptional role of Nub-PD during syncytial divisions.

### Dynamic subcellular localization of Nub-PD protein during mitosis

To further elucidate the role of Nub-PD during mitosis we followed its localization during the mitotic phases in *Drosophila* syncytial embryos. Nub immunostaining was very dynamic, with punctate staining in interphase nuclei (Fig. 6A). After nuclear envelope breakdown (NEB), at the onset of metaphase, Nub staining was restricted within the spindle bundles, more intense around the chromosomes, but not on centrosomes (Fig. 6A). At anaphase-telophase, Nub was recruited back to the newly formed nuclei and was cleared from the central spindle/midzone (Fig. 6A, C), indicating its active translocation during mitosis. To rule out fixation artifacts (Teves et al., 2016) we fixed embryos at low formaldehyde concentration (0.5%) and observed the same Nub enrichment within the spindle envelope (Schweizer et al., 2014) (Fig. S6A). Furthermore, live imaging of S2 cells transfected with constructs expressing mKO2- and mRFP-tagged Nub-PD also showed similar spindle enrichment during metaphase (Fig. 1H and Fig. S1H). Analyses of endogenously GFP-tagged Nub-PD in two independent knock-in fly lines (*sfGFP-nubPD*) and *PBac (nub-GFP)* using GFP antibodies, revealed identical localization pattern (Fig. 6B and Fig. S6B). In contrast, staining with an antibody against the related paralog Pdm2 (Fig. S6D-E), showed a different localization pattern. Although Pdm2 was present in interphase nuclei like Nub, it was absent from spindle microtubules during metaphase, but enriched in the midzone and centrosomes (Fig. S6C). These findings clearly demonstrate a dynamic and specific localization of Nub-PD during mitosis (Fig. 6C), consistent with its role in spindle organization and morphology in both syncytial embryos and S2 cells.

**Figure 6.**
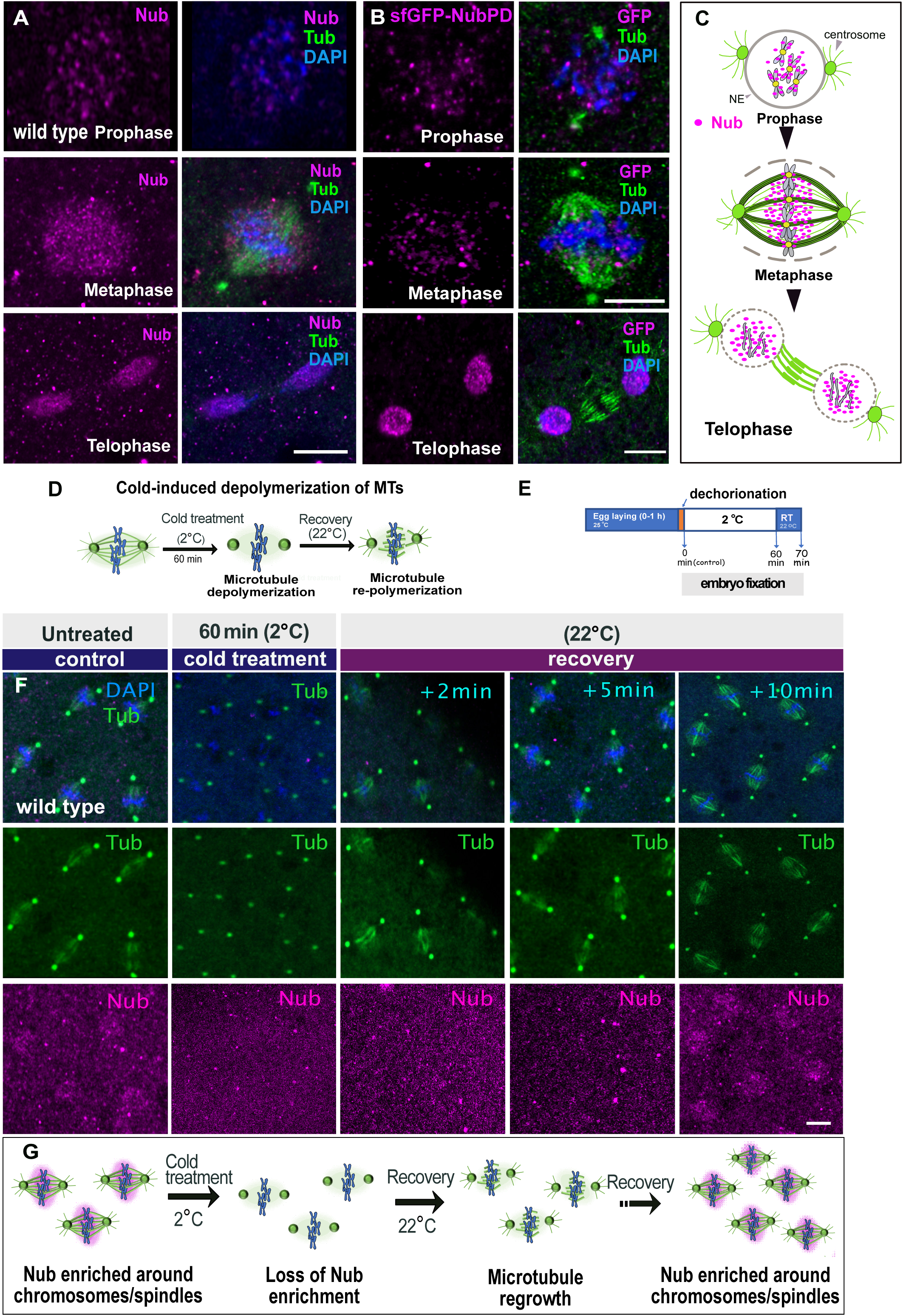
The dynamic localization of Nub within the mitotic spindle requires intact spindle microtubules. (A) Airy-scan confocal images (projections) of wild-type syncytial embryos showing the localization of Nub-PD (magenta), α- and γ-tubulin (Tub, green), and DNA (blue) during prophase, metaphase, and telophase. (B) Airy-scan Confocal projections showing *sfGFP-NubPD* syncytial embryos stained with antibodies for GFP (magenta), α- and γ-tubulin (Tub, green), and DNA with DAPI (blue) during prophase, metaphase, and telophase. (C) Graphical illustration describing the dynamic localization of Nub-PD (magenta) during prophase, metaphase and telophase. Centrosomes (green); nuclear envelope (NE, grey). (D) Schematic illustration of the spindle effects by cold-induced treatment during metaphase. Microtubules (MTs) depolymerized and spindles disassembled upon cold treatment, but centrosomes remained intact. (E) Diagram showing the procedure of cold-induced treatment in syncytial embryos. Embryos were shifted to room temperature up to 10 min to allow MT regrowth before fixation and staining. (F) Confocal images showing representative spindles of wild-type syncytial embryos showing the fluorescence signals of Nub-PD (magenta), α- and γ-tubulin (green), and DNA (blue) after 0 min (control) and 60 min of cold treatment. Post-treatment recovery of 2, 5, and 10 min is indicated. (G) Schematic illustration of Nub localization after cold treatment. The panel indicates the loss or alteration of Nub enrichment around the spindle when microtubules are depolymerized, illustrating its dependency on intact spindle microtubules.

### Nub-PD localization requires intact spindle-microtubules in an interdependent manner

To determine whether the Nub-PD enrichment at metaphase spindles depends on intact microtubules, we used non-invasive, cold-induced depolymerization of microtubules at the syncytial blastoderm stage (Hayward et al., 2014) (Fig. 6D-E). Incubation of wild type embryos at 2°C (60 min) completely eliminated tubulin staining at metaphase and also abolished Nub-PD enrichment around the spindles. Spindles quickly reformed upon recovery at room temperature, but Nub-PD did not reappear around the spindles until about 10 minutes, presumably at the following nuclear division (Fig. 6F-G). This indicates that Nub-PD localization relies on intact spindle microtubules and possibly other mitotic factors that support its spindle targeting.

The nuclear proteins Skeletor, Megator, East, and Chromator are localized throughout spindle microtubules by forming a matrix during the partially open mitosis of *Drosophila* syncytial embryos (Schweizer et al., 2014; Yao et al., 2018). This spindle matrix has been proposed to act as a mechanical element that mediates the interaction and movement of the kinetochore towards the poles (Qi et al., 2004; Zheng, 2010). To analyze a possible connection between Nub-PD and this matrix in maintaining spindle integrity, we carried out immunostaining of the spindle matrix protein Megator. Megator showed a similar localization pattern as Nub-PD in wild type embryos, and this did not change in *nub*^1^ mutant embryos (Fig. S6F). In contrast to Nub-PD, Megator remained intact in a mesh-like structure around the spindles after cold treatment (Fig. S6G), consistent with other studies using nocodazole as depolymerizing factor (Yao et al., 2018). Thus, Nub-PD but not Megator requires the presence of spindle microtubules for its localization during mitosis (Fig. 6F-G and Fig. S6G), indicating distinct localization requirements. In conclusion, our data argues against Nub-PD being an interacting partner within the matrix. Rather, our findings indicate that Nub-PD associates with microtubules to maintain spindle shape, length and integrity during the rapid syncytial nuclear divisions.

Next, we used the same assay to examine how efficient the *nub^1^* nuclei are in re-nucleating and re-stabilizing MTs once conditions become permissive again. We subjected wild type and *nub^1^* embryos to cold treatment and followed the microtubular regrowth at room temperature, either live or after fixation. In untreated embryos, microtubule spindle regrowth was visible within 2 minutes (Fig. 7A and S6H-I). On the contrary, cold-treated *nub^1^*embryos showed delayed microtubule regrowth during metaphase (Fig. 7A-C), and live imaging revealed that proper spindle architecture was not maintained at anaphase onset (Fig. 7D). Instead, abnormally elongated astral-like microtubules were formed, extending from the centrosomes (Fig. 7E).

**Figure 7.**
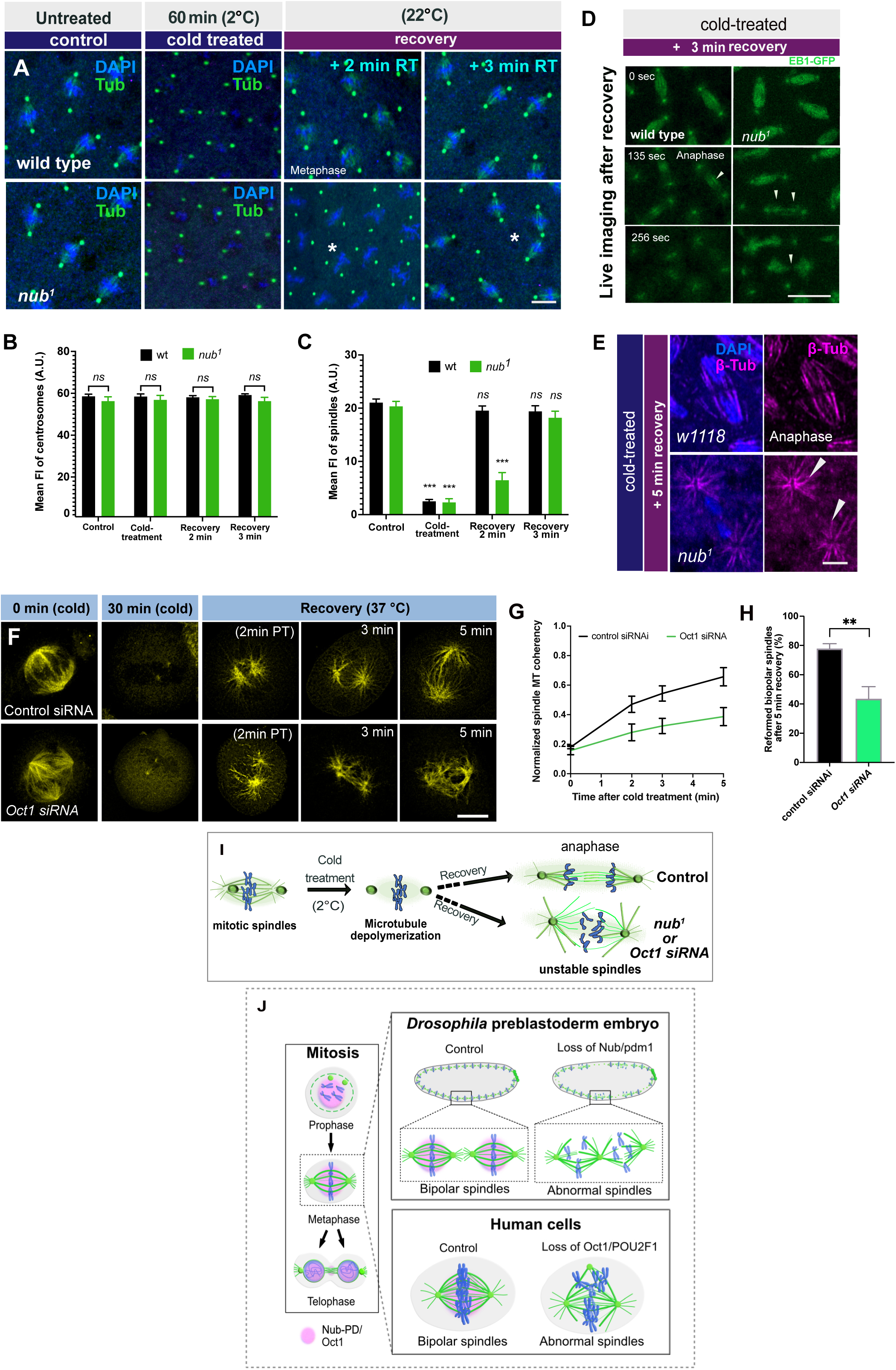
Nub-PD and POU2F1/Oct1 are required for spindle microtubule dynamics during reconstitution and regrowth. (A) Confocal images of representative spindles in wild-type and *nub^1^* embryos stained for α- and γ-tubulin (green). Embryos were cold-treated for 0 min (control) and 60 min. The recovery time is indicated (min). Scale bar, 10 μm (B–C) Quantification of centrosomal (B) and spindle (C) microtubule fluorescence intensity in wild-type and *nub¹* embryos under control conditions, cold treatment, and during MT regrowth (2–5 min recovery). *n* > 350 centrosomes and N > 5 embryos per condition. Error bars represent SEM. ns, not significant; **p < 0.01; p < 0.0001. (D) Selected confocal frames from live imaging of dividing wt and *nub^1^* nuclei after 3 min of recovery at room temperature following cold treatment. Arrows depict interpolar microtubules in the midzone during anaphase. (E) Confocal frames of dividing nuclei of wt and *nub^1^* during anaphase stained for β-tubulin (green) and DAPI (blue). Arrowheads indicate unstable spindle bundles of *nub^1^*embryos, forming aster-like structures. (F) Confocal images of representative mitotic spindles in live HeLa cells treated with control or POU2F1/Oct1-siRNA. Cells were incubated at 0.5–2D°C for 30 minutes (cold) and in ViaFluor® 488 to visualize spindle microtubules (yellow). Time points (in minutes) indicate post treatment (PT) recovery duration at 37°C during mitotic progression. (G–H) Normalized spindle MT coherency over time after cold treatment in control versus POU2F1/Oct1 siRNA–treated cells, showing impaired MT reassembly upon POU2F1/Oct1 knockdown (G). Spindle MT organization was quantified from α-tubulin images using the OrientationJ coherency index in Fiji/ImageJ, which reports local filament alignment (0 = random, 1 = perfectly ordered) (see Material and Methods for details). Percentage of cells that reformed bipolar spindles 5 min after recovery, demonstrating a significant reduction in POU2F1/Oct1-depleted cells (H). *p* < 0.001. mean ± SEM, (values are a sum of 3 independent experiments, ≥ 60 cells per condition). (I) A schematic representation showing the microtubule regrowth in wild type and *nub^1^* or POU2F1/Oct1-siRNA after cold treatment. Microtubule regrowth was aberrant after cold treatment in both *nub^1^* mutants and POU2F1/Oct1-siRNA knockdown, leading to unstable spindle formation. (J) Schematic summary of the observed phenotypes showing microtubule regrowth defects in syncytial *Drosophila* embryos and human cells. Loss of Nub/Pdm1 leads to unstable microtubule bundles within the spindle, resulting in misorientation of the division axis and aberrant interactions with neighboring spindles. In human cells, downregulation of POU2F1/Oct1 disrupts spindle organization and bipolarity, leading to the formation of extra spindle poles.

Notably, the loss of Nub-PD disrupted the organization of the interpolar microtubule (iMTs) array in the midzone, leading to unstable spindles (Fig. 7D). These results further support the role of Nub-PD in maintaining MT stability in spindles, enabling them to resist poleward forces during anaphase. Additionally, a portion of *nub^1^* cold-treated embryos (30%) failed to reform spindle microtubules (Fig. S6H-I), suggesting a possible contribution of Nub-PD for efficient spindle microtubule re-nucleation after induced depolymerization. Taken together, our data argue that Nub-PD is important for both the stability and reformation of the mitotic spindle, with implications for microtubule nucleation and organization during mitosis.

Next, we investigated whether human POU2F1/Oct1 plays a similar role in spindle reconstitution and regrowth. Using the same cold treatment assay (for 30 minutes), we found a defect in spindle reassembly upon POU2F1/Oct1 depletion. In control cells, bipolar spindles rapidly reformed within a few minutes after temperature recovery, whereas POU2F1/Oct1-depleted cells showed delayed and disorganized spindle regrowth (Fig. 7F). Quantification confirmed a significantly reduced spindle reformation rate and lower spindle coherency in POU2F1/Oct1-knockdown cells compared to controls (Fig. 7G,H). These results demonstrate that POU2F1/Oct1 is required for efficient spindle microtubule regrowth following depolymerization.

Together, these findings suggest that Nub-PD and POU2F1/Oct1 may share common functions in maintaining stable spindle microtubule dynamics during mitosis, when microtubules are continuously reorganized (Fig. 7I-J).

### Factors affecting Nub-PD recruitment to mitotic spindles

To identify the molecular interactions controlling the dynamic recruitment of Nub-PD during nuclear division cycles, we carried-out a targeted maternal RNAi-knockdown screen of components of the mitotic spindle machinery during syncytial nuclear divisions, followed by Nub immunostaining. We analyzed 29 genes, using 45 independent RNAi or mutant fly lines, and out of these, nine genes affected the localization of Nub during mitosis (Table S1). Several of these genes specifically altered the distribution of Nub-PD during M-phase but not during S-phase (interphase) (Fig. S7B, C) and this pattern was consistent between early and late NCs (Table S1). We consider the effects of these latter genes to be more specific and likely to reflect a direct role in regulation of Nub localization, rather than secondary consequences of general cell cycle disruption. More specifically, downregulation of the kinases Nek2 and Niki/Nek9 drastically reduced Nub-PD localization specifically in metaphase, suggesting phosphorylation by these kinases is essential for the translocation of Nub-PD to the spindles (Fig. S7A-C). Similarly, knockdown of Klp61F (the ortholog of vertebrate Kinesin-5/EG5) and Klp3A abrogated Nub-PD localization in metaphase but not in interphase (Fig 8D and S7A-C). The expression level of Klp61F in syncytial mitosis is controlled by the Crb, Galla-2 and Xpd (CGX) complex (Hwang et al., 2020). Our analysis showed that RNAi of Crb, but not of Galla-2 or Xpd, affects Nub-PD localization around the spindles (Fig. S6A and Table S1). Crb, Klp61F are localized in spindles and exhibited mitotic phenotypes similar those seen in *nub^1^* mutants during syncytial divisions (Hwang et al., 2020).

**Figure 8.**
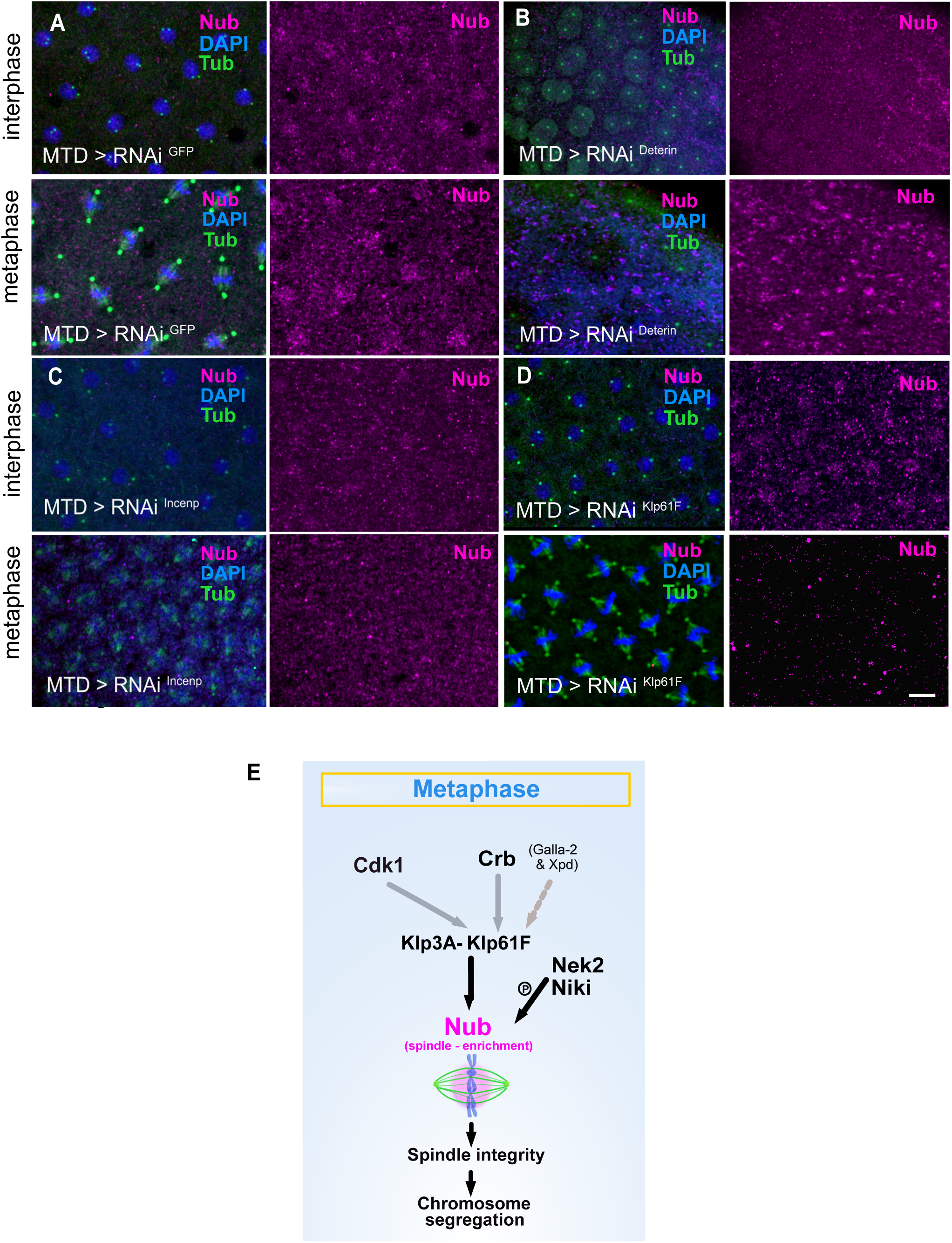
Factors affecting Nub-PD recruitment to mitotic spindles. (A-D) Airy-scan confocal images displaying the cortex of *MTD>RNAi^GFP^ (control) (A)*, *MTD>RNAi^Deterin^* (B), *MTD>RNAi^Incenp^* (C) and *MTD>RNAi^Klf61F^* (D) syncytial embryos stained with antibodies for Nub (magenta), β + γ-tubulin (green) and DAPI (blue). Mitotic phases of nuclei are indicated (prophase, metaphase). (E) Proposed model of Nub-PD recruitment to the mitotic spindle, incorporating findings from both current (black) and previously published work (grey). The recruitment of Nub-PD to the mitotic spindle involves mitosis-related motor proteins Klp61F and Klp3A, the polarity determinant Crb and the kinases Cdk1, Nek2, and NiKi/Nek9. The enrichment of Nub around the spindle supports spindle integrity and accurate chromosome segregation.

The CPC is a highly conserved master regulator of mitosis that is required for spindle and kinetochore assembly. In mammalian cells, it is composed of four subunits: the enzymatic component Aurora B (AurB) and the three regulatory and targeting components Incenp, Survivin and Borealin (Carmena et al., 2012). Our analysis shows that knockdown of AurB, Incenp and of the Borealin-related protein Deterin/Survivin abolished Nub-PD staining in interphase/prophase nuclei and metaphase spindles (Fig. 8A-C, S7A-C and Table S1). Further, Cdk1 disruption affected Nub-PD localization both in interphase and metaphase, indicating that Nub-PD localization, and presumably its function, is downstream of the mitotic-entry signals (Table S1). Because knockdowns of AurB, Incenp, and Deterin/Survivin caused strong mitotic and developmental defects (Table S1), we cannot rule out that the severe mis-localization of Nub in these embryos (interphase and metaphase) is a secondary effect, due to the overall mitotic or developmental phenotypes caused by depletion of these mitotic components. However, downregulation of Crb, Klp3A, Klp61F, Nek2, Niki caused reduced Nub-PD signal within metaphase spindles, but not in interphase, and overall milder spindle mitotic defects under the RNAi conditions. In these RNAi experiments the reduction of Nub-PD levels was only partial (Table S1), and did not appear to fall below a threshold sufficient to disrupt mitotic progression. This is consistent with our observations in *nub^1^/+* heterozygous embryos (Fig. S5A), where Nub-PD signal is reduced but mitosis proceeds normally. Taken together, our work identifies a regulated and precisely timed translocation of Nub-PD from the chromatin to the spindle envelope during the rapid nuclear cycles in the syncytial embryo.

## Discussion

Nub/Pdm1 has been shown to be required for proliferation in the *Drosophila* embryonic nervous system, wing discs and midgut epithelium (Cifuentes and García-Bellido, 1997; Doe, 2017; Ng et al., 1995; Tang et al., 2018; Tran and Doe, 2008; Tsuji et al., 2008). However, how Nub is controlling proliferation during development was not known and its role in mitosis had not been explored. Several transcription and splicing factors have been suggested to confer direct mitotic functions based on their localization to the mitotic apparatus, but very few have been confirmed experimentally by loss-of-function analyses (Pellacani et al., 2018; Somma et al., 2020). Here, we show that the Nub-PD isoform is not just a transcriptional regulator in interphase but also a critical mitotic factor during early embryogenesis. We demonstrated that it is required for timely progression of the rapid and synchronous syncytial nuclear-divisions by maintaining mitotic spindle integrity and elongation. Importantly, this function is independent of its transcriptional activity, revealing a previously unrecognized, non-transcriptional role for Nub-PD in early development.

Using live imaging in cells and syncytial embryos, we show that Nub-PD localizes to within the spindle envelope. This enrichment depends on intact microtubule and likely regulated by mitotic factors, including the Cdk1, Nek2, and NiKi/Nek9 kinases, the motor proteins Klp61F and Klp3A, and the polarity determinant Crb. Similar regulatory mechanisms have been reported for human POU/Oct proteins, including Nek6 phosphorylation of POU2F1/Oct1 for spindle pole localization in human cells (Kang et al., 2011) and AurB/Cdk1 phosphorylation of POU5F1/Oct4 during G2/M to regulate its chromatin dissociation in embryonic stem cells (Kim et al., 2018; Shin et al., 2016). Interestingly, we found that sequence-specific DNA binding and transcriptional regulation is not important for Nub-PD localization to the spindle, and likely not for its mitotic functions as we did not observe any mitotic defects. In addition, the closely related Pdm2 paralog, which contains almost identical DNA binding POU and homeo domains (98% and 79% identity respectively), did not predominantly localize to mitotic spindles.

Except for these domains, Nub and Pdm2 do not show regions of high amino acid identity. Taken together, these findings and the mitotic phenotypes observed in transcriptionally inactive *nub^1^, MTD>nub^RNAi^* and *nub^1^;mat>nub^RNAi^* embryos, strongly support that Nub-PD plays a critical non-transcriptional function in mitosis of early embryonic development, in addition to its role in transcriptional regulation later during development. Similarly, our findings suggest that the human POU2F1/Oct1 confers important roles in mitotic progression. Thus, Nub-PD and POU2F1/Oct1 play dual roles, functioning as transcriptional regulators during interphase and in differentiated cells, and as key contributors to mitotic progression in actively dividing cells.

Understanding exactly how Nub-PD is involved in the direct regulation of mitotic progression is challenging. Our data show that both centrosome numbers and morphology remain unaffected, and instead, centrosomes frequently detached from the mitotic spindles as a consequence of spindle collapse. Despite being detached, these free centrosomes retain immunoreactivity for γ-tubulin and the capacity to duplicate, indicating that centrosome formation and integrity does not involve Nub-PD. Instead, data from syncytial embryos support a model in which Nub-PD interacts, directly or indirectly, with spindle microtubules, rather than centrosomal microtubules or the spindle matrix, through a mutually dependent interaction. Intact microtubules and the identified mitotic proteins are required to ensure proper localization of Nub-PD within the spindles, and thereby its function in supporting proper spindle architecture and dynamics during metaphase and anaphase. Consistent with this, cold-depolymerization and recovery reveals that *nub^1^* embryos can initiate MT regrowth around chromosomes, but fail to convert these regrowing MTs into a stable bipolar spindle, suggesting inefficient coupling and integration of chromosome-derived MTs with centrosome-derived arrays leading to disorganized spindles and progressively defected midzone bundles. Together, our data suggests a model in which Nub-PD maintains spindle stability by promoting the organization and stabilization of spindle and mid zone MTs. In a subset of embryos, the complete absence of MT regrowth is consistent with a role for Nub-PD in supporting efficient spindle microtubule regrowth, either by influencing early nucleation events at γ-tubulin–containing sites or by stabilizing nascent α/β-tubulin polymers during spindle reassembly. Given that Nub is a chromatin-associated DNA-binding transcription factor, we also cannot exclude a contribution via Ran- and CPC-dependent chromatin-based microtubule nucleation, although this remains speculative at present. Nub-PD may be anchored in the spindle apparatus via other MT-associated proteins (MAPs), not identified here. One possible model is that Nub-PD, together with MAPs, promotes the organization of microtubule bundles, including the sliding of antiparallel microtubules to ensure proper spindle elongation. Recent work has shown that Klp61F physically interacts with Crb, Galla-2, and Xpd (CGX complex) in syncytial embryos (Hwang et al., 2020). However, how the CGX complex and each component regulates Klp61F levels and activity is unclear. Loss of Nub-PD in the *nub^1^* mutant causes defects in nuclear syncytial divisions, identical to those caused by Crb depletion (Hwang et al., 2020). In addition, knockdown of Crb, but not Galla-2 or Xpd, disrupts Nub’s enrichment around the spindle. These results suggest that Nub acts downstream or in parallel with Crb and the motor protein Klp61F to promote proper spindle architecture and elongation during anaphase (Fig. 8E).

It is interesting to note that many MAPs involved in spindle assembly undergo liquid–liquid phase separation and form condensates on microtubules (Sun et al., 2024). Importantly, both immunostaining and live imaging revealed that Nub-PD is present in punctate structures within the spindles (Fig. 1H, 6A, and Fig. S1M), raising the question if Nub-PD functions by acting within condensates on microtubules. Protein prediction tools for unstructured regions (IUPred2A)(Erdos and Dosztanyi, 2020; Meszaros et al., 2018) and for three-dimensional structures (AlphaFold2)(Jumper et al., 2021; Varadi et al., 2024) show that outside the DNA binding domains, Nub-PD is highly disordered, except for one coiled-coil structure (not shown).

Interestingly, POU2F1/Oct also contains a coiled-coil structure in an otherwise primarily unstructured protein, besides the DNA binding domains. The exact molecular mechanism how Nub-PD promotes anaphase spindle stability and proper elongation, along with structure-function analyses, will need further investigation.

Many human POU/Oct proteins are linked to cancer development or progression, and high expression of POU proteins in malignant cells is correlated with poor patient prognosis(Ben-Batalla et al., 2010; Castrillo et al., 1991; Jullien et al., 2015; Rudin et al., 2019; Vázquez-Arreguín et al., 2019; Vázquez-Arreguín and Tantin, 2016). In fact, POU2F1/Oct1 is expressed in all human tumor cell lines and it has been found to be proto-oncogenic in epithelial malignancies(Jafek et al., 2019; Stepchenko et al., 2022; Vázquez-Arreguín et al., 2019; Vázquez-Arreguín and Tantin, 2016). However, the molecular role of POU2F1/Oct1 in tumor proliferation remains unclear. We show in live HeLa and HEK293 cells that POU2F1/Oct1 is crucial for the normal progression of mitosis and that its downregulation results in spindle collapse leading to prolonged mitotic progression, and the formation of multinucleated cells. These results correlate with earlier findings in fixed HeLa cells, showing that both up- and downregulation of POU2F1/Oct1 caused abnormal mitosis (Kang et al., 2011). The phenotypes we observe upon POU2F1/Oct1 depletion may be explained by distinct mechanisms, in which centriole amplification or fragmentation could drive spindle multipolarity, while cytokinesis failure could give rise to multinucleation. Both scenarios remain possible, and additional experiments will be required to distinguish between centrosome-related defects and cytokinesis failure in this context. Our live imaging in a cold-induced microtubule regrowth assay demonstrates that POU2F1/Oct1 is required for maintaining spindle architecture and promoting rapid spindle microtubule regrowth following microtubule depolymerization.

This phenotype is remarkably similar to that observed upon loss of the *Drosophila* Nub-PD isoform, supporting a similar and potentially conserved function of POU/Oct factors in stabilizing spindle microtubule organization. This points at new roles for POU/Oct proteins in cellular proliferation and tumorigenesis, however, the molecular mechanism underlying this mitotic function and its broader relevance for tumor progression remain to be elucidated. Whether this mitotic role of POU2F1/Oct1 is mechanistically distinct from its transcriptional activity also remains to be determined.

Our results suggest an important contribution of POU2F1/Oct1 and Nub-PD in efficient spindle organization and chromosome segregation in systems with fast divisions, such as tumor cells and syncytial embryos. These so far overseen, additional roles of POU/Oct proteins in mitosis, may explain some of the unanswered questions regarding their oncogenic properties, and open new possibilities in drug targeting and pharmaceutical interventions to block cell proliferation and malignant growth.

## EXPERIMENTAL MODEL AND STUDY PARTICIPANT DETAILS

### Drosophila melanogaster

Flies were grown on standard cornmeal/yeast food at 25°C, and experiments were performed at 25°C unless otherwise specified. *Drosophila melanogaster* stocks used in this study and associated references are listed in the key resources Table S2. Some stocks were treated with tetracycline for two to three generations to be cleaned from *Wolbachia* infection, which then was verified by PCR.

## METHOD DETAILS

### Drosophila Genetics

Details for the source of these genotypes are in the key resources Table S2. CyO and TM3 balancer strains carrying GFP transgenes were used to identify embryos with the desired genotypes for fly genetics.

### Germline-specific RNAi knockdown

Transgenic flies carrying RNAi vectors (Valium20 or Valium22), specifically designed for optimal expression of RNAi hairpins in the germline (Blake et al., 2017) were used. Homozygous males from the RNAi lines were mated with virgin females of the Maternal Triple Driver (MTD)-Gal4, which drives expression throughout oogenesis. *MTD>shRNA* females from this cross were backcrossed to the homozygous *UAS-shRNA* or to F1 *MTD>shRNA* males. Their F2 embryo progenies were analyzed.

### Nub-PD degradation in the early embryo using deGradFP system

To degrade Nub-PD protein in early embryos, *nos>deGradFP* (Gaskill et al., 2021) and *hb>deGradFP* (Vazquez-Pianzola et al., 2022) transgenic lines were used. Briefly, *nos>deGradFP* flies were crossed with *sfGFP-nubPD* and offsprings were selected for nos>*deGradFP/sfGFP-nub-PD*, and intercrossed. Similarly, *hb>deGradFP* flies crossed with *sfGFP-nubPD* and offsprings were selected for *sfGFP-nubPD*; *hb>deGradFP,* and intercrossed. From these crosses, embryos (0-2hr) were collected on agar plates supplemented with yeast at 25°C. Embryos from *nos>deGradFP* and *hb>deGradFP* females were used as a control.

### *D. melanogaster* cell culture and RNA interference

*D. melanogaster* Schneider’s (S2) cells, and S2 cells that stably express Histone 2B-GFP (H2B-GFP) and α*-*tubulin-mCherry were maintained in Schneider’s insect medium (GIBCO), supplemented with 10% heat-inactivated fetal bovine serum (FBS, GIBCO) and 1% Penicillin-streptomycin antibiotics cocktail, at 25°C without CO_2_.

Double-stranded RNA (dsRNA) targeting firefly luciferase (*Luc*) (Control) and *nub* transcripts (*nub-RB* and *nub-RD*) were synthesized using a dsRNA synthesis kit according to the manufacturer’s instructions. S2 cells were incubated with respective dsRNA for 48-72 hrs at 25°C.

### Human cell culture and small interfering RNAs (siRNAs)

HeLa cells were maintained in DMEM F12 or DMEM GlutaMax (GIBCO) media supplemented with 10% heat-inactivated FBS (GIBCO) at 37°C with 5% CO2. Cells were transfected with POU2F1/Oct1 siRNAs, targeting three independent regions of human POU2F1/Oct1 (sc-36119) and Control siRNA (sc-37007) (Santa Cruz Biotech), at 100nM siRNA concentrations. Transfections were carried out with lipofectamine-RNAiMAX reagent (Life Technologies) following the manufacturer’s instructions. Human cells were tested for mycoplasma contamination (IDEXX BioAnalytics) and were negative throughout the study.

### Molecular Biology and Transgenics

#### Generation of *sfGFP-nubPB* and *sfGFP-nubPD* flies by CRISPR/Cas9-mediated homology-directed repair (HDR)

##### Design and construction of gRNA-plasmids

We used the CRISPR optimal target finder algorithm (https://flycrispr.org/target-finder/) to identify suitable genomic target sequences in the first protein-coding exons of *nub-RB* (*nub* exon 2) and *nub-RD* (*nub* exon 4). The following CRISPR genomic target sites were selected for targeting Nubbin isoforms with GFP :

Nub-RB.Trg1 GGATGCTGAAACGCCTTCGGTGG

Nub-RB.Trg2 CGGATGATGATACAGCAGTGTGG

Nub-RD.Trg1 TGGTTATGTCGGAGCTACGTTGG

Nub-RD.Trg2 TTGGCACACCGCTAGTCCCGAGG

(PAM sequences are highlighted in bold). Target sequence-specific sense and antisense oligonucleotides (Star Methods/Resource Table S2) with 5’ complementary overhanging sequences to the BbsI-cut *pCFD3*-*dU6:3gRNA* vector were PNK-phosphorylated and annealed prior to ligation into BbsI-cut *pCFD3*-*dU6:3gRNA* (http://crisprflydesign.org/protocols/). The resulting guide plasmids were confirmed by Sanger sequencing (Eurofins Genomics).

##### Construction of pBS II SK(+)-Nub-RD.sfGFP and pBS II SK(+)-Nub-RB.sfGFP donor vectors

Approximately 1 kb long DNA sequences upstream and downstream of the genomic target sites were PCR-amplified from fly genomic DNA (BL#58492) with the Q5 DNA polymerase kit (NEB, # M0491S) using Nub-PD.HL.Fw and Nub-PD.HL.Rw, or Nub-PD.HR.Fw and Nub-PD.HR.Rw primers that added flanking restriction sites for *XhoI* and *HindIII* (Nub-PD.HL), or *HindIII* and *BamHI* (Nub-PD.HR) to the PCR products. XhoI/HindIII-digested Nub-PD.HL, and HindIII/BamHI-digested Nub-PD.HR were ligated in the XhoI/BamHI-cut *pBluescript* (referred to as *pBS*) *II SK(+)* vector. Nub-PDsfGFP.Fw and Nub-PDsfGFP.Rw primers were used to PCR-amplify the sfGFP-coding sequence from the *pScarlessHD-sfGFP-DsRed* vector (Addgene #80811), simultaneously adding 25 bp flanking sequences that overlapped with the Nub-PD.HL/Nub-PD.HR-hinge region.

The resulting PCR product was inserted into HindIII-linearized *pBSII SK(+)*-*Nub-PD.HL/Nub-PD.HR* with the HiFi DNA assembly kit (NEB, #E2621S). Similarly, we constructed *pBS II SK(+)-Nub-RB*.sfGFP using ∼1kb homology-providing sequences flanking the genomic target sites that were PCR-amplified from fly genomic DNA (BL#58492) with the primers Nub-PB.HL.Fw and Nub-PB.HL.Rw, or Nub-PB.HR.Fw and Nub-PB.HR.Rw adding flanking *XhoI*/*HindIII* or *BamHI*/*NotI* restriction sites. XhoI/HindIII-, respectively BamHI/NotI-cut, PCR-products were ligated into the respectively digested *pBS II SK(+)* vector. Nub-RB.sfGFPFw and Nub-RB.sfGFPRw primers were used to amplify the *sfGFP* CDS from *pScarlessHD-sfGFP-DsRed* simultaneously adding 25 bp flanking sequences that overlapped with the Nub-PB.HL/Nub-PB.HR-hinge region. The *sfGFP*-coding fragment was inserted into EcoRV/BamHI-opened *pBS II SK(+)*-*Nub-PB.HL/Nub-PB.HR* via HiFi DNA assembly (NEB). The final *pBS II SK(+)-Nub-RD.sfGFP* and *pBS II SK(+)*-*Nub-RB.sfGFP* donor constructs were verified by Sanger sequencing (Eurofins Genomics).

##### Fly injections and candidate screening

The Cas9-expressing fly line *y1,M[Act5C-Cas9]ZH-2A, w^1118^, DNAlig4^169^* (BL#58492) was used for embryo injections. A mix of two *pCFD3*-dU6:3gRNA guide vectors and the respective donor construct was co-injected into fly embryos. Each resulting G_0_ individual was crossed to *w^1118^* flies, and subsequent screening for GFP expression was done in the adult G_1_ population. GFP-positive G_1_ individuals were crossed to a *CyO* balancer stock, and the presence and correct integration of sfGFP were further confirmed by PCR on isolated genomic DNA. The Cas9 carrying allele *y1, M[Act5C-Cas]}ZH-2A, w^1118^, DNAlig4^169^* located on the first chromosome was segregated away. Homozygous *nub^sfGFP.Nub-PB^* and *nub ^sfGFP.Nub-PD^* fly stocks could be established and were further verified by genomic DNA sequencing (Eurofins Genomics), and analyzed for viability, life span, and fecundity. In *nub^sfGFP.Nub-PB^* flies, the DNA sequence coding for Val^2^-Lys^239^ of sfGFP is inserted between 2L:12,594,048 and 2L:12,594,049 [+], resulting in an n-terminal insertion of sfGFP-tag following Thr^17^ of the predicted Nub-PB protein. In the case of *nub^sfGFP.Nub-PD^* flies, sfGFP (Val^2^-Lys^239^) is inserted n-terminally after Leu^6^ of the predicted Nub-PD isoform (between 2L:12,619,013 and 2L:12,619,014 [+]). In both cases, the sfGFP coding sequence contains a short c-terminal extension (coding for Ile-Gln-Pro-Arg-Lys-Ile-Ile) originally present in the *pScarlessHD-sfGFP-DsRed* plasmid.

### Antibody production and Immunostaining

Antibodies against Pdm2-PA/PB isoforms were raised in rabbits against a synthesized peptide (PPKRLAEEQEEEK) conjugated to keyhole limpet hemocyanine carrier protein (Thermo Fisher Scientific, Waltham, MA, USA). The Pdm2 peptide without the carrier protein was coupled to cyanogen bromide-activated Sepharose 4B according the manufacturer’s protocol (Sigma-Aldrich, St Louis, MI, USA), and used for affinity-purification of the antisera.

#### Embryo

Embryos were bleached, dechorionated, and fixed for 20 minutes in 4% formaldehyde (or for 15 min in 0.5% formaldehyde) and incubated overnight at 4°C with primary antibodies. Secondary antibodies conjugated to Cy3, Cy5, Alexa Fluor-488 or Alexa Fluor-568 were diluted as recommended by the manufacturers and embryos were incubated for 2 hours at room temperature. Stained embryos were mounted in a Vectashield Plus antifade mounting medium.

#### S2 and HeLa cells

Cells were seeded and fixed in 4% formaldehyde for 30 min. Fixed cells were blocked in PBS-Triton X100 (PBST) supplemented with 0.5% Normal goat serum (NGS) for 60 min at room temperature and incubated with primary antibody overnight at 4°C. The next day, cells were washed and incubated with a secondary antibody for 2hr at room temperature. DNA was counterstained with DAPI. Cells were mounted in 90% glycerol for imaging.

### Injections of anti-Nub and Imaging

Dechorionated embryos were placed on coverslips coated by a heptane/glue mix and then covered with halocarbon oil. Monoclonal mouse anti-Nub (Nub 2D4) was diluted in 6% dimethyl sulfoxide (DMSO) in deionized H_2_O. Nub antibody (12 μM) and 3 KDa Alexa Fluor™ 488 Dextran (10mg/ml) were injected into embryos using a microinjection system (FemtoJet, Eppendorf) coupled to an inverted fluorescence microscope (Cell Observer, Zeiss). Injections were done at the anterior part of the embryo. The injected embryos were immediately placed to an AxioImagerZ1 (Zeiss) microscope for live imaging by using a 20x/0.75NA Plan-APOCHROMAT objective (Carl Zeiss).

### Live Imaging

#### Embryo

Embryos (0-1h old) were dechorionated for 3 min in 7% sodium hypochlorite solution and mounted onto a pre-prepared slide (Tsarouhas et al., 2019). For wide-field live imaging, embryos were imaged with a CCD camera (AxioCam 702, Carl Zeiss) attached to an AxioImagerZ1 (Zeiss) microscope by using a 20x/0.75NA Plan-APOCHROMAT objective (Carl Zeiss). Individual Z-stacks with a step size of 1.2–1.9 μm were taken every 1 or 2 min over a 2–6-h period. Embryos were collected from yeasted grape juice agar plates aged at 25°C. For confocal imaging, embryos were imaged with a laser-scanning confocal microscope (LSM 780, Carl Zeiss) using a 63 × Plan-APOCHROMAT /1.4 M27 Oil-immersion objective. For high-resolution confocal imaging, an airy-scan-equipped confocal microscope system (Zeiss LSM 800, Carl Zeiss) with a 63 × Plan-APOCHROMAT /1.4 M27 oil-immersion objective was used. Raw data were processed with the airy-scan processing tool of the Zen Black software version 2.3 (Carl Zeiss). Images were converted into tiff format using ImageJ/Fiji software (http://rsb.info.nih.gov/ij/).

#### S2 and HeLa cells

For analyzing the mKO2- and RFP-tagged Nub-PD protein localization, S2 cells were seeded into 35 mm glass bottom cultured plates (MatTek) overnight. Cells were transfected with 100 µg of pW8 (carrier plasmid) using Effectene transfection kit (Qiagen). After 48hr of transfection, cells were treated with ViaFluor 488 for 20 min to visualize the microtubules. Images were taken every 1 minute for a period of 2-4 hrs. For mitotic phenotypes, S2 cells that stably expressed Histone 2B-GFP (H2B-GFP) and α-tubulin-mCherry were treated with control or *nub-RD* dsRNA for 72 hr. For cell imaging, Zeiss LSM 800 Airy-scan confocal microscope with a Plan/Apo 63 x 1.4 NA oil immersion objective was used. Images were analyzed using Zeiss 2011 (Blue) software and were exported in tiff format. The tiff files were imported into ImageJ/Fiji to generate the videos.

### RNA extraction and RT-qPCR

Embryos were collected for 2hr, subsequently washed and dechorionated. Syncytial blastoderm-staged embryos were collected in Trizol. S2 cells were harvested from 6-well plates, washed with PBS, and resuspended in triazole. RNA extraction and RT-qPCR were performed as previously described (Lindberg et al., 2018).

### Cold-Induced MT Depolymerization

For cold-treatment assays, embryos (0-2 hrs) were collected, dechorionated, placed on ice-cold Petri dishes, and incubated on ice inside Polystyrene-boxes at 4°C incubator room. Following 60 min treatment on ice, embryos were fixed at 0 sec (immediately after cold treatment and inside the cold room), 60 sec, 120 sec, 180 sec, 5 min or 10min after shifting them to 25°C. For live imaging, dechorionated embryos were mounted onto a slide with a gas permeable membrane and covered with halocarbon oil (#700, Sigma)(Tsarouhas et al., 2019). Slides were placed on ice-cold Petri dishes and incubated on ice for 60 min, before quick transfer to the microscope (within 30-60 sec) and imaged immediately, focusing on the centrosomes (Conduit et al., 2015; Hayward et al., 2014). Similarly, human cells growing in cultures glass bottom dishes were incubated on ice for 30 min inside Polystyrene-boxes at 4°C incubator room. The treatment duration was optimized in pilot experiments to induce spindle microtubule depolymerization in over 85% of cases without impairing the capacity for regrowth. A digital thermometer, equipped with probes attached inside the dishes, continuously recorded the temperature, which was constantly kept at 2.0 ± 0.5 °C.

## QUANTIFICATIONS AND STATISTICAL ANALYSIS

### Quantifications of nuclei dynamics

Confocal micrographs were transformed into the “binary” images followed by the Fiji/ImageJ plugin “Segmentation Editor” image for threshold setting and automatic calculation. The relative nuclear to cytoplasmic ratio (N/C ratio) was calculated on binary images as: *N/C = E_nu_/(E_cyto_ = E _total_ - E_nu_),* where *E_nu_* = area of nuclei, E_cyto_= area of the cytoplasm and *E_total_*=area of the whole embryo (the visible part). *E_nu_* was defined by the His2A-RFP or DAPI signals after manually tracing the borders of nuclei. *E_total_* was estimated by the fluorescence signal of the whole embryo. Cortex area with nuclei gaps was calculated after tracking the cortical area without nuclei (gaps) and dividing this area by the area of the whole embryo.

For the time profile of mitotic phase in nuclei of syncytial embryos, S phase is defined as the period from early interphase to the late prophase, when chromosomes just begin to condense and become visible. M phase includes the time of metaphase (chromosomes in the equator), anaphase until telophase.

### Quantifications of RFI and spindle dynamics

Image processing and quantification were performed using the Fiji. Confocal images were exported in *tiff* format (uncompressed) by Zeiss 2011 (Blue) software. Relative fluorescence intensity (RFI) of Nub was calculated by dividing the average fluorescence intensity of Nub signals in the nucleus or spindles (FIn or FIs) by the fluorescence intensity of the cytoplasm (FIc) outside the nucleus or the spindle area. *RFI = FI_n_/ FI_c_ or FI_s_/ FI_c_*. The fluorescence intensity was measured using a region of interest (ROI) defined by the limits of the nuclei in prophase (DAPI signal) or by the spindles in metaphase (Tubulin signal). The background was measured with the equivalent ROI outside the area of the nuclei or the spindle. Images were sum-intensity Z projections.

The pole-to-pole distance is the length of a line (μ*m*) drawn from one spindle pole to the other. For the pole-to-pole distance as a fraction of time (μ*m/sec*), the length of the line between the spindle poles was calculated in each time point based on EB1-GFP real-time imaging data. The quantification was performed using Fiji and the “Profile module” of Zen2011.

### Quantifications of centrosome fluorescence intensity and density

Centrosome density was determined by counting centrosomes within 50 × 50 µm regions of interest and normalizing to the ROI area (centrosomes/µm^2^). For quantification of centrosome intensity, maximum-intensity projections were generated and individual centrosomes were segmented as diffraction-limited spots using Fiji. For each centrosome, a fixed-size circular region of interest (ROI; corresponding to 0.5–0.7 µm diameter) was placed over the centrosome signal, and the mean fluorescence intensity was measured after subtraction of the local cytoplasmic background (measured in a nearby ROI lacking centrosome signal).

### Quantifications of MT organization

To quantify spindle microtubule organization (related to Figure 7G), we analyzed α-tubulin fluorescence using the OrientationJ plugin (v3.0, BIG-EPFL, Fiji). Maximum-intensity projections from images were background-subtracted (rolling-ball radius 30–50 px) using identical settings for all samples. A fixed spindle region of interest (ROI) was defined manually for each cell and used for all corresponding timepoints. OrientationJ was run in Structure Tensor mode. The plugin produced a coherency image in which pixel intensities (0–1) represent the local degree of filament alignment. For each frame, the mean coherency value within the spindle ROI was measured using Analyze and Measure tool. Result tables were exported as CSV files for subsequent plotting in GraphPad Prism.

### Statistics

Error bars are standard error of means ± S.E.M. Graphs with box-plots show the median (horizontal line) and the range from 25th to 75th percentile. The bars depict maximum and minimum values. *p*-values were calculated through the *t*-test for unpaired variables if the data passed normality test (Shapiro-Wilk test). For grouped data, one-way or two-way ANOVA followed by Tukey’s *post hoc* test for multiple comparisons was applied. For spindles dynamics (Fig. 3H and 3I), data analysis was performed using JMP Pro 17.0 statistical program (SAS Institute). Normality was determined by the D’Agostino-Pearson normality test, using a significance cut-off value of *p* = 0.05. The *n* values of the quantifications are provided in the figure legends. All plots and statistical tests were performed on GraphPad Prism 10.1.

## STAR * Methods

### RESOURCE AVAILABILITY

#### Lead Contact

Further information and requests for resources and reagents should be directed to and will be fulfilled by the lead contact, Ylva Engström.

#### Materials Availability

Fly-lines and plasmids generated in this study are available upon request from the lead contact.

#### Data and code availability

- All original data reported in this paper will be shared by the lead contact upon request.
- This paper does not report any original code.
- Any additional information required to analyze the data reported in this paper is available from the lead contact upon request.

#### Author contributions

Conceptualization, P.G, V.T. and Y.E.; Methodology, Investigation, and Formal analysis, P.G, V.T, L.K, S.S; Visualization P.G, V.T, L.K, Writing-Review & Editing, P.G., V.T. and Y.E.; Funding Acquisition, Y.E. and V.T.

## Acknowledgments

Stocks obtained from the Bloomington Drosophila Stock Center (NIH P40OD018537) were used in this study. A monoclonal Nub antibody used in this study was developed by Michalis Averof and Steve Cohen, and was obtained from the Developmental Studies Hybridoma Bank, created by the NICHD of the NIH and maintained at The University of Iowa, Department of Biology, Iowa City, IA 52242. We thank Cai Yu, Xiaohang Yang, and William Chia for the Pdm1/Nub antibody, and Georg Wolfstetter for critically reading and commenting on the material and methods section. The authors also acknowledge the technical support from the IFSU (Imaging Facility at Stockholm University). This work was financially supported by The Swedish Cancer Society (20 1044 PjF and 23 2963 Pj) to Y.E, The Swedish Research Council (2018-04401) to Y.E. and by O. E. och Edla Johanssons vetenskapliga stiftelse and Magn. Bergvalls stiftelse to VT.

