## Supplementary Figures and Supplementary Tables for "A Non-Transcriptional Mitotic Function of POU/Oct Factors Ensures Spindle Stability and Chromosome Segregation"

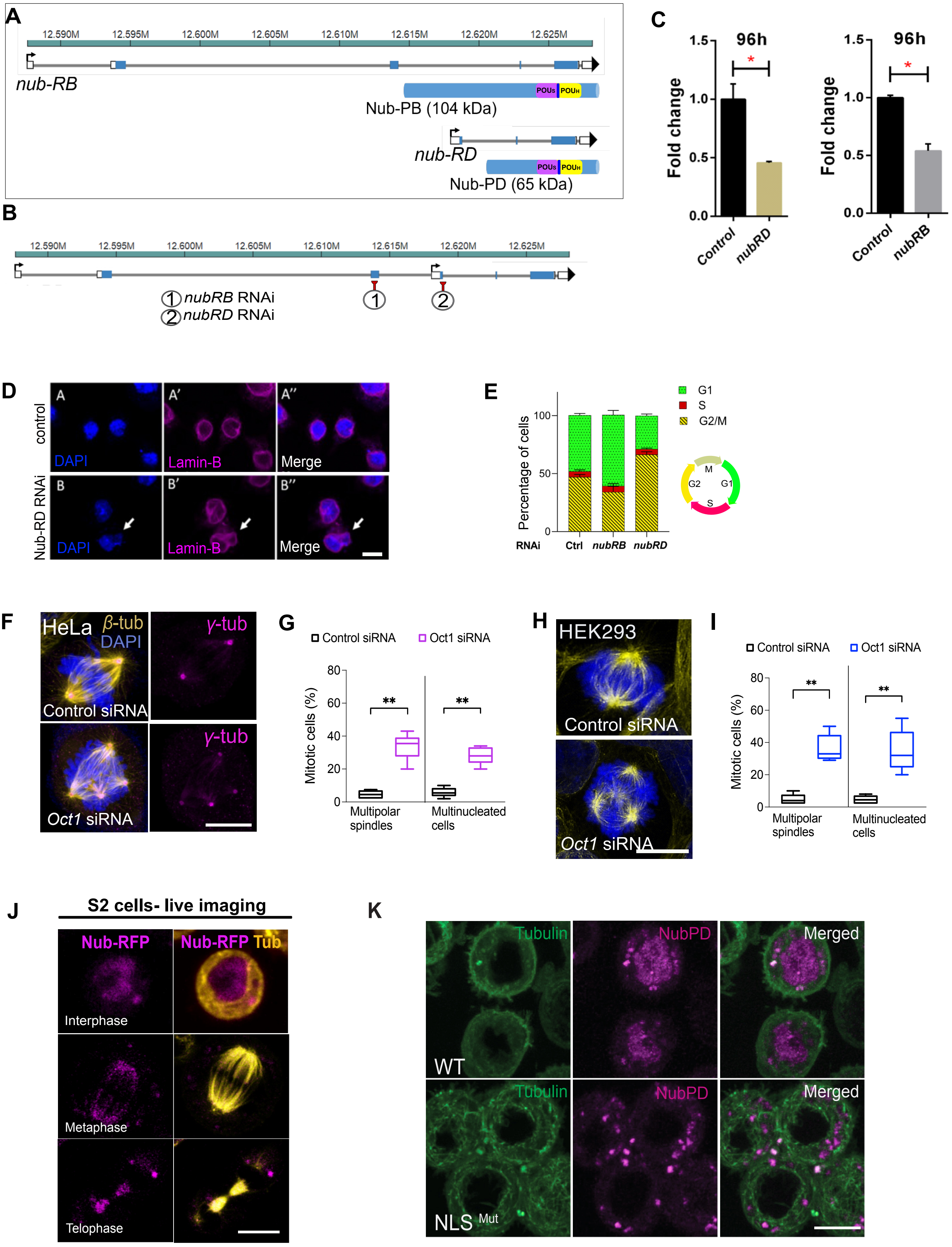
A
nub-RB
Nub-PB (104 kDa)
nub-RD
Nub-PD (65 kDa)
B
nubRB RNAi
nubRD RNAi
C
96h
Fold change
Control
nubRD
nubRB
D
control
Nub-RD RNAi
DAPI
Lamin-B
Merge
E
Percentage of cells
G1
S
G2/M
RNAi
Ctrl
nubRB
nubRD
F
HeLa
β-tub
DAPI
γ-tub
Control siRNA
Oct1 siRNA
G
Control siRNA
Oct1 siRNA
Mitotic cells (%)
Multipolar spindles
Multinucleated cells
H
HEK293
Control siRNA
Oct1 siRNA
I
Control siRNA
Oct1 siRNA
Mitotic cells (%)
Multipolar spindles
Multinucleated cells
J
S2 cells- live imaging
Nub-RFP
Nub-RFP Tub
Interphase
Metaphase
Telophase
K
Tubulin
NubPD
Merged
WT
NLS Mut

Figure S1

**Figure S1. RNAi-mediated depletion of *nub* transcripts in S2 cells. Spindles defects in POU2F1/Oct1-depleted HeLa and HEK293 cells.**

(A) Schematic illustration of the *nub* gene and its transcripts (*nub-RB* and *nub-RD*) initiated at two independent transcription start sites (arrows), encoding two protein isoforms, Nub-PB (104 kDa) and Nub-PD (65 kDa) respectively. Solid blue boxes indicate coding exons. The location of the POU-specific domain (POUs) (magenta) and POU-homeodomain ($POU_H$) (yellow), is indicated.
(B) Schematic illustration of *dsRNAs* specifically targeting the *nub-RB* transcript (exon 3) (1) and the *nub-RD* transcript (exon 1 of this transcript and exon 4 in the *nub* gene, not present in the *nub-RB* transcript) (2).
(C) Measurement of *nub-RB* and *nub-RD* transcripts by RT-qPCR to confirm reduced levels of the respective transcript. The bar graph represents the fold change ($log^2$). * $p < 0.05$; Statistical significance was calculated by students *t*-test.
(D) Representative images of control (A–A″) and *nub-RD* dsRNA-treated S2 cells (B–B″) stained with DAPI and Lamin-B antibody. Arrows indicate multinucleated cells observed upon Nub depletion.
(E) Stacked box graphs representing the distribution of cell cycle phases in control (*luc*-RNAi), *nub-RB* RNAi and *nub-RD* RNAi-treated $R^+$ FUCCI S2-cells.
(F) Confocal images of mitotic HeLa cells (metaphase) treated with control-siRNA or POU2F1/Oct1-siRNA and processed for immunofluorescence with β-tubulin (yellow), and γ-tubulin (magenta). DNA was counterstained with DAPI (blue).
(G) Quantification of mitotic defects (multipolar spindles and multiple nuclei) in control-siRNA and POU2F1/Oct1-siRNA-treated HeLa cells. Statistical significance was calculated by Mann Whitney test. ** $p < 0.01$, ($N$ = 6 individual experiments and $n$ = 490 mitotic cells).
(H) Confocal images of mitotic HEK293 cells (metaphase) treated with control-siRNA and POU2F1/Oct1-siRNA and processed for immunofluorescence with β-tubulin and γ-tubulin (yellow). DNA was stained with DAPI (blue).
(I) Quantification of mitotic defects (multipolar spindles and multiple nuclei) in control-siRNA and POU2F1/Oct1-siRNA-treated HEK293 cells. Statistical significance was calculated by Mann Whitney test. ** $p < 0.01$, ($N$ = 3 individual experiments and $n$ = 149 mitotic cells).
(J) Subcellular localization of Nub-PD-RFP protein (magenta) in mitotic S2 live-cells stained with the membrane-permeable ViaFluor 488 dye to visualize the spindle microtubules (yellow). Representative images of Nub-PD-RFP localization during interphase, metaphase and telophase are shown.
(K) Confocal image-projections of live S2 cells transfected with Nub-PD $^{WT}$-mKO2 or NubPD $NLS^{Mut}$-mKO2. Cells were incubated with ViaFluor® 488 dye to visualize the microtubules (green). Mutations in the NLS of Nub-PD-mKO2 blocked its nuclear localization in interphase cells.

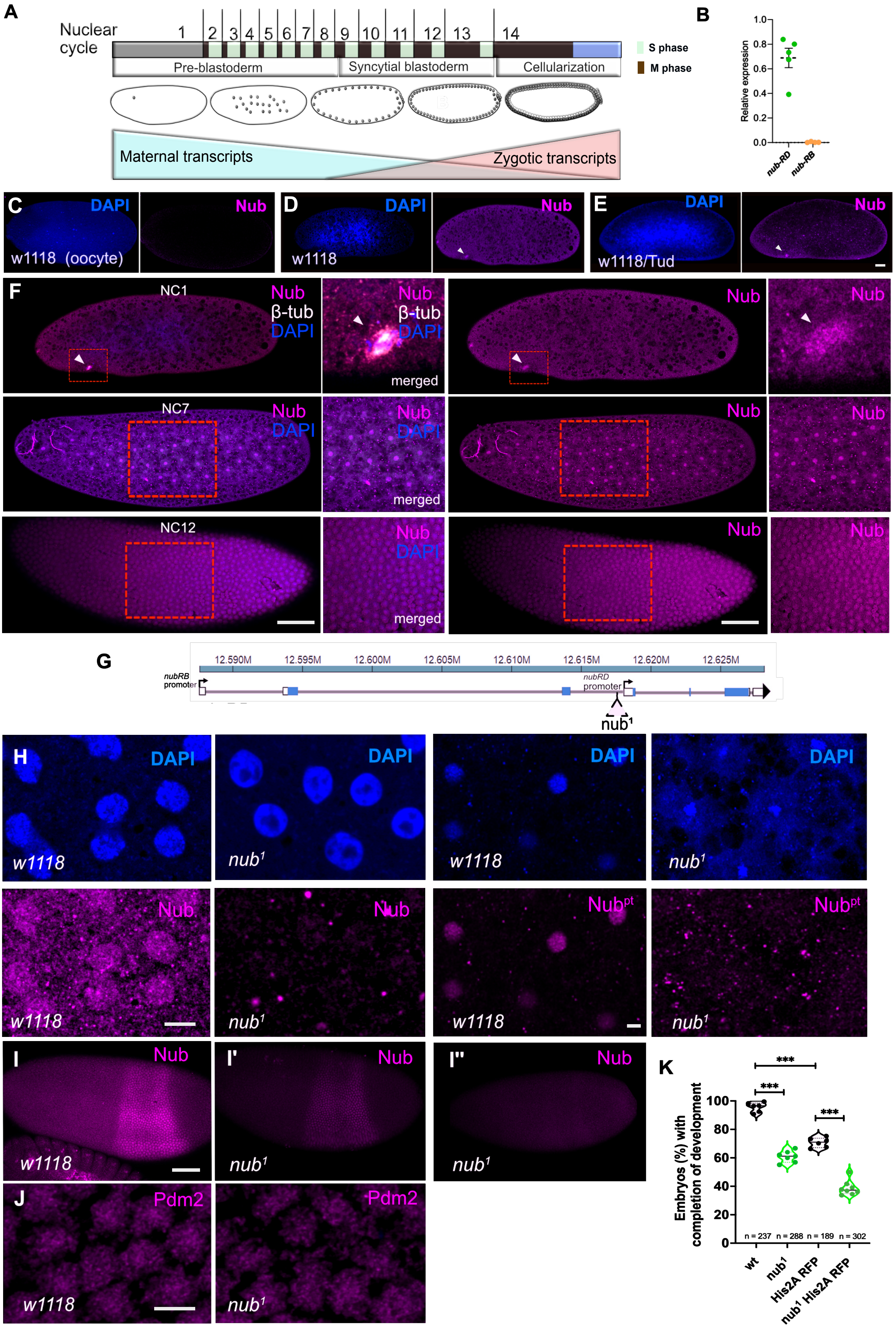
A
Nuclear cycle
1 2 3 4 5 6 7 8 9 10 11 12 13 14
Pre-blastoderm
Syncytial blastoderm
Cellularization
S phase
M phase
Maternal transcripts
Zygotic transcripts
B
Relative expression
nub-RD
nub-RB
C
DAPI
Nub
w1118 (oocyte)
D
w1118
E
w1118/Tud
F
NC1
Nub
β-tub
DAPI
merged
NC7
NC12
G
nubRB promoter
nubRD promoter
nub1
H
w1118
nub1
Nubpt
I
I'
I''
J
Pdm2
K
Embryos (%) with completion of development
n = 237
n = 288
n = 189
n = 302
wt
nub1
His2A RFP
nub1 His2A RFP
***

Figure S2

**Figure S2. Nub-PD expression during the syncytial stage.**

(A) Schematic drawing showing the early nuclear divisions and cortical migration during *Drosophila* embryogenesis, and the profile of maternal and zygotic transcripts (Kwasnieski et al., 2019). Nuclear Cycle (NC) 1 is initiated after the fusion of the male and female pro-nuclei. In NC 4–6, nuclei divide and spread out along the anterior–posterior axis (axial expansion). During NC 8–10, nuclei migrate to the cortex of the embryo (cortical migration). At the posterior end, pole cells (germ cells) are formed (cycle 9). At NC 10-13, nuclei divide at the cortex. At NC 14, membranes invaginate from the surface and enclose each nucleus, forming the cellular blastoderm. NCs are rapid and synchronous and switch between S and M phases. Maternally provided transcripts are predominant, and no transcription occurs before NC8. Zygotic transcripts can be measured for a few genes from NC8 and then transcription increases gradually during the following nuclear division cycles (Kwasnieski et al., 2019).
(B) RT-qPCR analysis of the relative transcript levels of *nub* (*nub-RD* and *nub-RB*) in 0-2h embryos.
(C-E) Late oocytes (stage 14) (C) and syncytial embryos (NC1) from respective genotypes (D-E) stained for antibodies against Nub protein (magenta) and DNA with DAPI (blue).
(F) Confocal images showing whole mount syncytial embryos at NC1, NC7 and NC12 stained with anti-Nub (magenta), anti-β-tubulin (white) and DAPI (blue). The arrowheads denote the mitotic spindles at NC1. Images on the right column are zoomed-in areas of the embryo cortex indicated by the rectangular frames in the left column images.
(G) Schematic drawing showing the insertion of the 412 retrotransposon in the promoter region near the transcription start of *nub-RD*, and in the intronic region of the *nub-RB* transcript.
(H) Syncytial embryos of $w^{1118}$ and $nub^1$ stained with anti-Nub antibody or anti-Nub peptide ($Nub^{pt}$) (magenta) and DAPI (blue).
(I-I'') Confocal images showing whole syncytial $w^{1118}$ and $nub^1$ embryos at NC14 stained with anti-Nub (magenta). The signal of Nub is either partially (I') or severely lost (I'') in $nub^1$ embryos.
(J) Confocal images showing syncytial $w^{1118}$ and $nub^1$ embryos at NC11 prophase stained with anti-Pdm2 (magenta). The signal of Pdm2 protein is not affected in $nub^1$ embryos.
(K) Graphs showing the percentage of wild type ($n$ = 237), $nub^1$ ($n$ = 288), His2A-RFP ($n$ = 189) and $nub^1$ His2A-RFP ($n$ = 302) embryos with completed embryonic development. Values are means of 6 or 7 repeated experiments. Statistical significance was calculated by unpaired two-tailed $t$ -test, *, $p < 0.03$, **, $p < 0.01$, ***, $p < 0.001$. Error bars show s.e.m. Scale bars, 50 μm (b, c) and 10 μm (e-g).

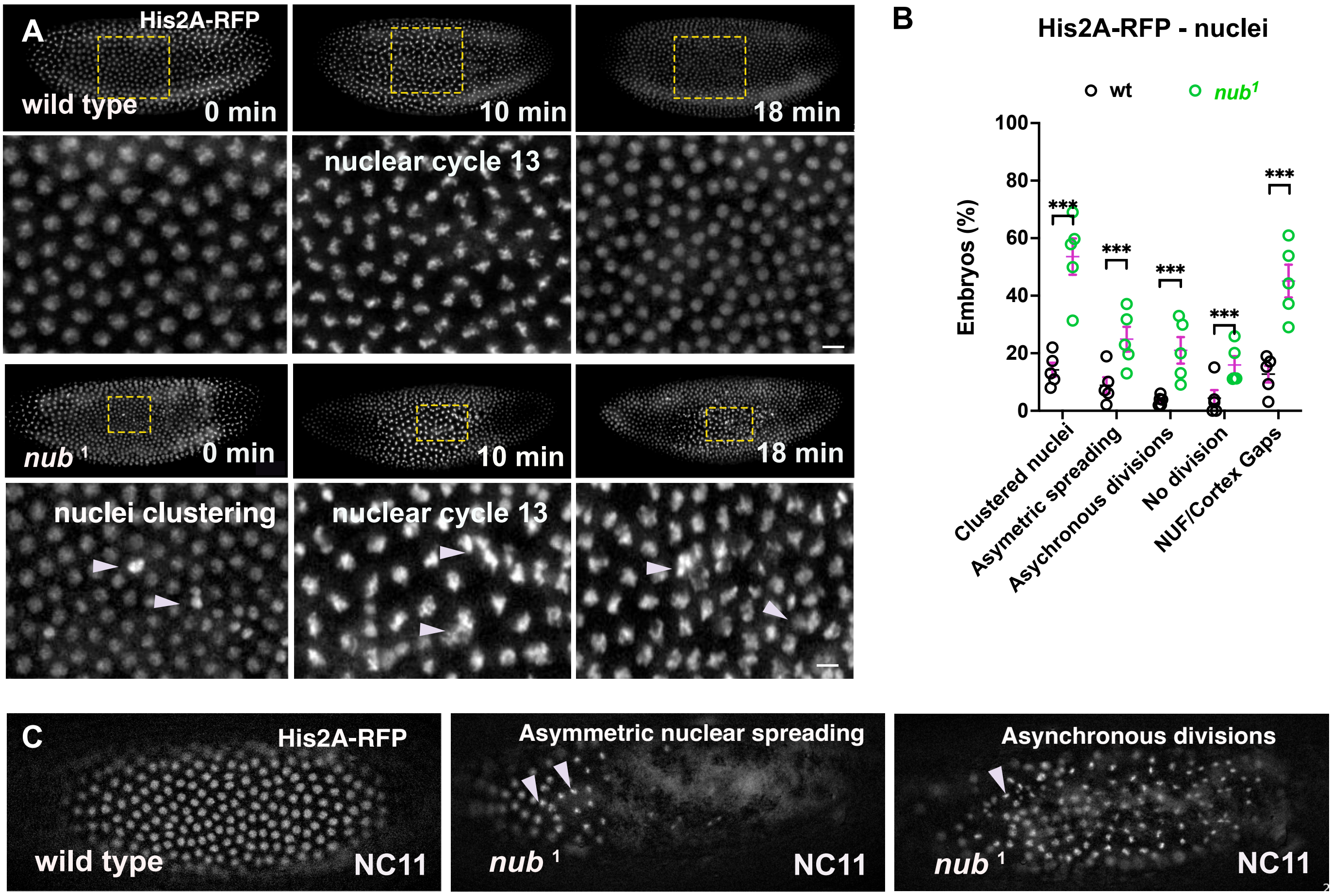

**Figure S3. Phenotypic analysis of nub1 mutant in syncytial nuclear divisions**

(A) Fluorescent images from time lapse recordings showing live wild type and *nub¹* mutant embryos at NC13. Embryos are expressing the His2A-RFP reporter. Images below depict zoomed-in areas of the embryo cortex indicated by the rectangular frames in the upper images.
(B) Graph shows the percentage of wild type (n = 48) and *nub¹* (n = 86) embryos with clustered, asymmetric, asynchronous, arrested nuclei and nuclear fall-out phenotypes. Values represent the means of five repeated experiments.
(C) Fluorescent images from time-lapse recordings showing live wild type and *nub¹* mutant embryos at NC11, expressing the nuclear His2A-RFP reporter. *nub¹* mutant embryos show asymmetrically distributed nuclei and asynchronous divisions (arrow heads). Scale bars 10 µm.

Figure S3

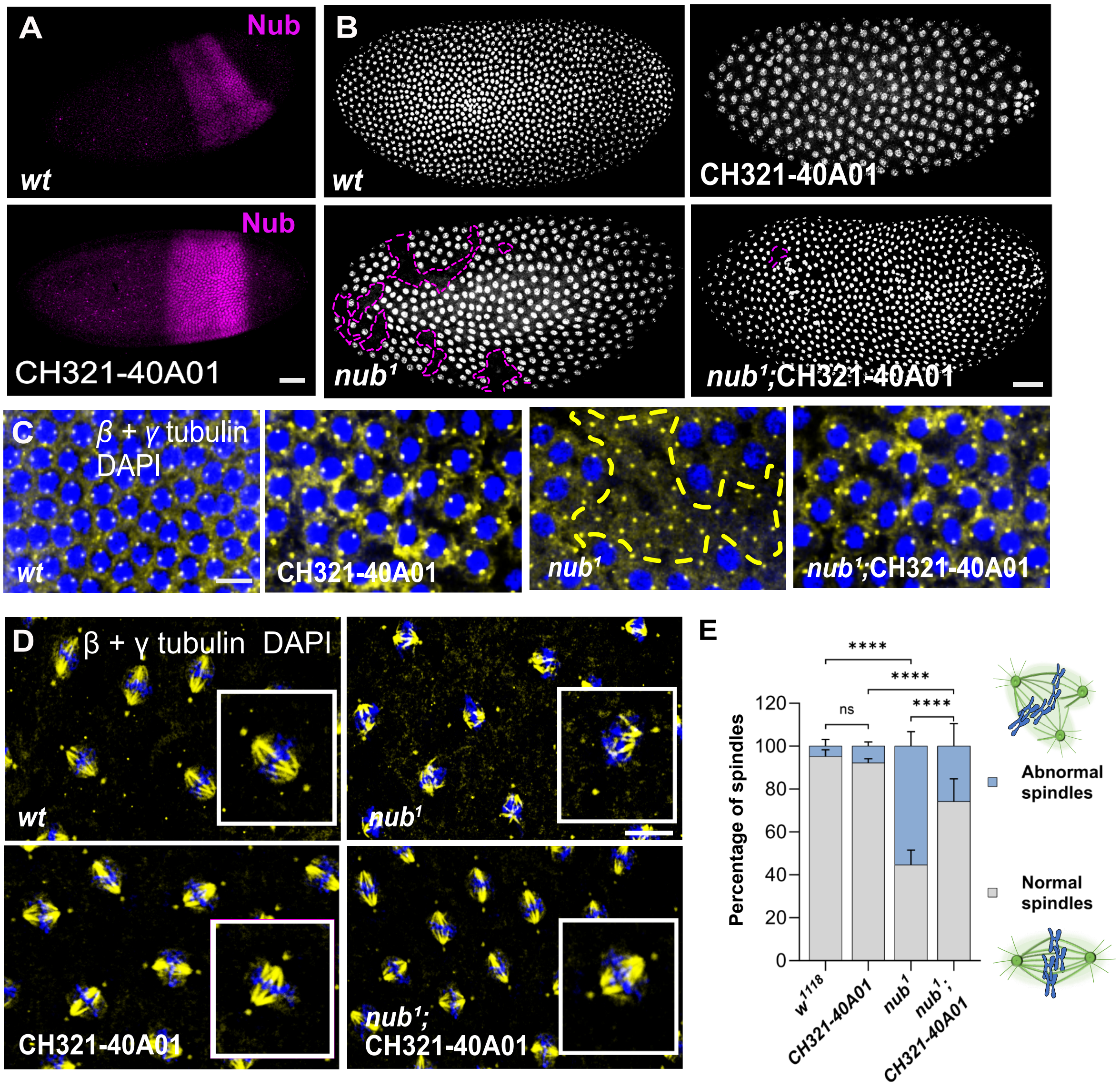

**Figure S4. A nub duplication-locus rescues the mitotic defects of *nub¹* mutant.**

(A) Representative images of embryos after zygotic genome activation, when the maternally expressed nub RNA and protein is degraded, and a broad band of newly transcribed and translated Nub-PD is visible, by incubation with anti-Nub in wt and CH321-40A01 embryos.
(B) Representative images of mitotic nuclei in syncytial embryos of the indicated genotypes: $w^{1118}$, CH321-40A01, $nub^1$ and $nub^1$; CH321-40A01 (rescue genotype).
(C-D) Confocal images of syncytial embryos stained for β -tubulin, γ- tubulin (yellow) and DNA with DAPI (blue). The genotypes of w1118, CH321-40A01, $nub^1$ and $nub^1$; CH321-40A01 (rescue genotype) are shown during NC12 (C) or NC8 (D). Insets are zoomed-in areas of the corresponding image. Dashed lines indicate nuclei-free gaps and areas with free centrosomes on the cortex of $nub^1$ embryos.
(E) Quantification of normal and abnormal spindles in syncytial embryos of the indicated genotypes (NC 6-10). n = 30-50 mitotic spindles, N =10 syncytial embryos. Statistical significance was assessed by two-way ANOVA followed by Fisher's multiple comparison test. **** p < 0.0001, * p<0.05, ns = not significant.

**Figure S4**

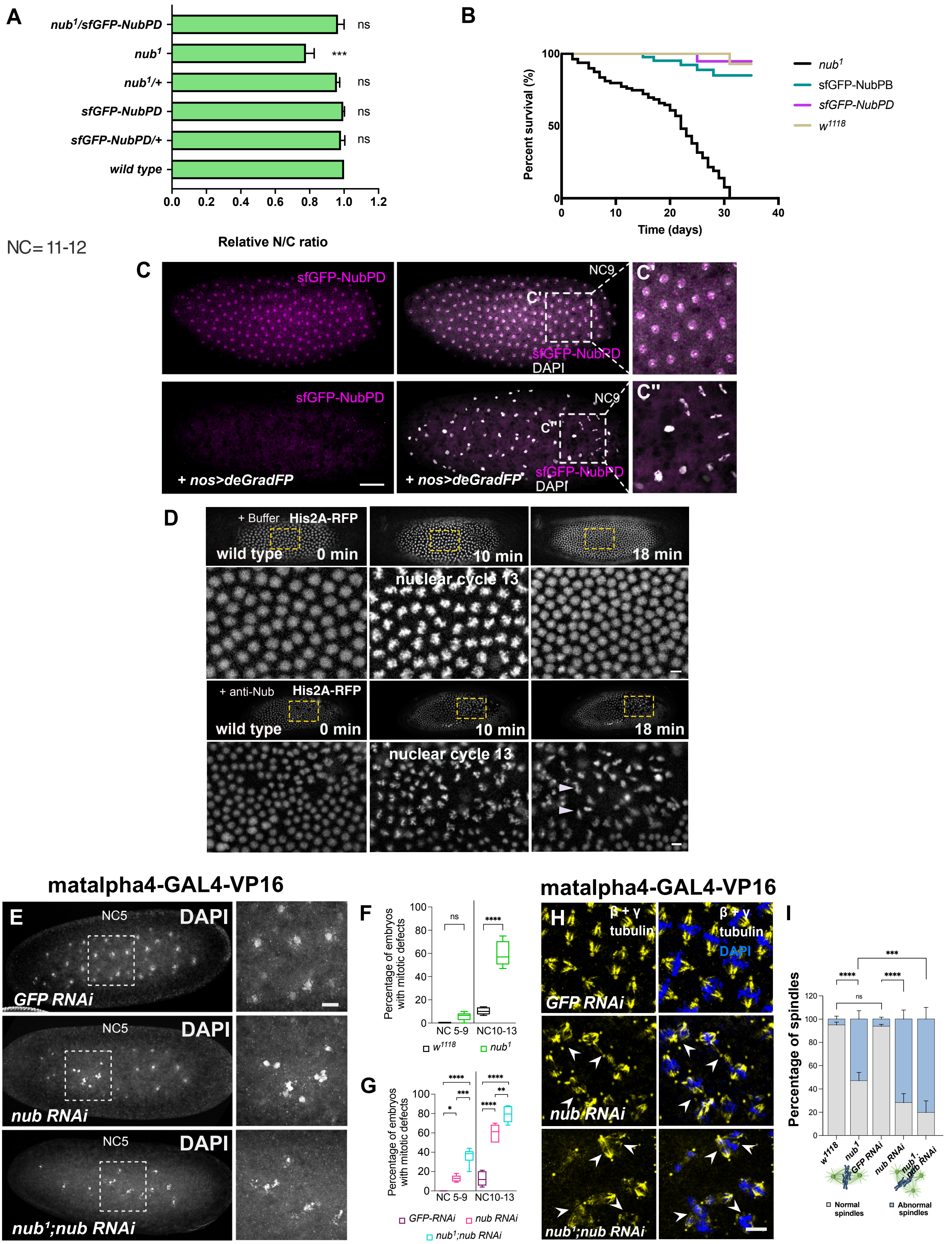
A
nub1/sfGFP-NubPD
nub1
nub1/+
sfGFP-NubPD
sfGFP-NubPD/+
wild type
ns
***
0.0 0.2 0.4 0.6 0.8 1.0 1.2
Relative N/C ratio
NC= 11-12
B
Percent survival (%)
Time (days)
nub1
sfGFP-NubPB
sfGFP-NubPD
w1118
C
sfGFP-NubPD
DAPI
NC9
C'
C''
+ nos>deGradFP
D
+ Buffer
His2A-RFP
wild type
0 min
10 min
18 min
nuclear cycle 13
+ anti-Nub
matalpha4-GAL4-VP16
E
NC5
DAPI
GFP RNAi
nub RNAi
nub1;nub RNAi
F
Percentage of embryos with mitotic defects
NC 5-9
NC10-13
G
GFP-RNAi
H
β + γ tubulin
I
Percentage of spindles
Normal spindles
Abnormal spindles

Figure S5

**Figure S5. Nub-PD loss leads to severe mitotic defects as early as NC5.**

(A) Quantification of relative nuclear-to-cytoplasmic (N/C) ratio across indicated embryo genotypes during NC11–12. ns, not significant; ***p < 0.001 as calculated by unpaired two-tailed *t* -test.
(B) Survival curves of adult flies carrying *sfGFP-NubPD, sfGFP-NubPB, nub*$^{1}$ *and w1118* genotypes. No major differences in lifespan are observed between *sfGFP-NubPD* and wild type.
(C) Fluorescent images of control (*sfGFP-NubPD*) and of *sfGFP-NubPD; nos>deGradFP* embryos in NC9 stained for DNA with DAPI (white), showing loss of GFP (magenta) in *sfGFP-NubPD; nos>deGradFP* embryos. Images on the right (C', C'') depict zoomed-in areas of the embryo cortex indicated by the rectangular frames in images on the left.
(D) Fluorescent images from live imaging showing live wild type embryos injected with buffer and anti-Nub antibody. Embryos are expressing the His2A-RFP reporter. Images below indicate zoomed-in areas of the embryo cortex indicated by the rectangular frames in the upper images. Arrowheads indicate abnormal nuclei divisions in anti-Nub antibody injected embryos. Nuclei in embryos injected with buffer were intact.
(D) Graph shows the percentage of wild-type embryos injected with buffer *(n* = 28*)* and anti-Nub *(n* = 37*)* antibody with attached, asymmetric, asynchronous, arrested nuclei and nuclear fall-out (NUF) phenotypes. Values represent means of five repeated experiments.
(E) Representative images of syncytial embryos with respective genotypes stained with DAPI (white). *GFP-RNAi* (P[matalpha4-GAL4- VP16]> GFP-RNAi), *nub-RNAi* (P[matalpha4-GAL4-VP16]> *nub*-RNAi), *nub*$^{1}$; *nub-RNAi* (*nub*$^{1}$; P[matalpha4-GAL4- VP16]> *nub*-RNAi). Images on the right are zoomed-in areas indicated by the rectangular frames in images on the left.
(F) Quantification of the percentage of embryos with mitotic defects in *w*$^{1118}$ (control) and *nub*$^{1}$ embryos at NC5-9 and NC10-13. *** denotes statistical significance *p < 0.0001,* calculated by unpaired two tailed *t* -tests.
(G) Quantifications showing the percentage of embryos with mitotic defects in GFP-RNAi (control), *nub-RNAi,* and *nub*$^{1}$;*nub-RNAi* conditions. The mitotic defects were scored during NC5-9 and NC10-13. *, *p <0.05,***, *p <0.01,* ***, *p <0.0001*. Statistical significance was calculated by one-way ANOVA followed by Tukey's multiple comparison test.

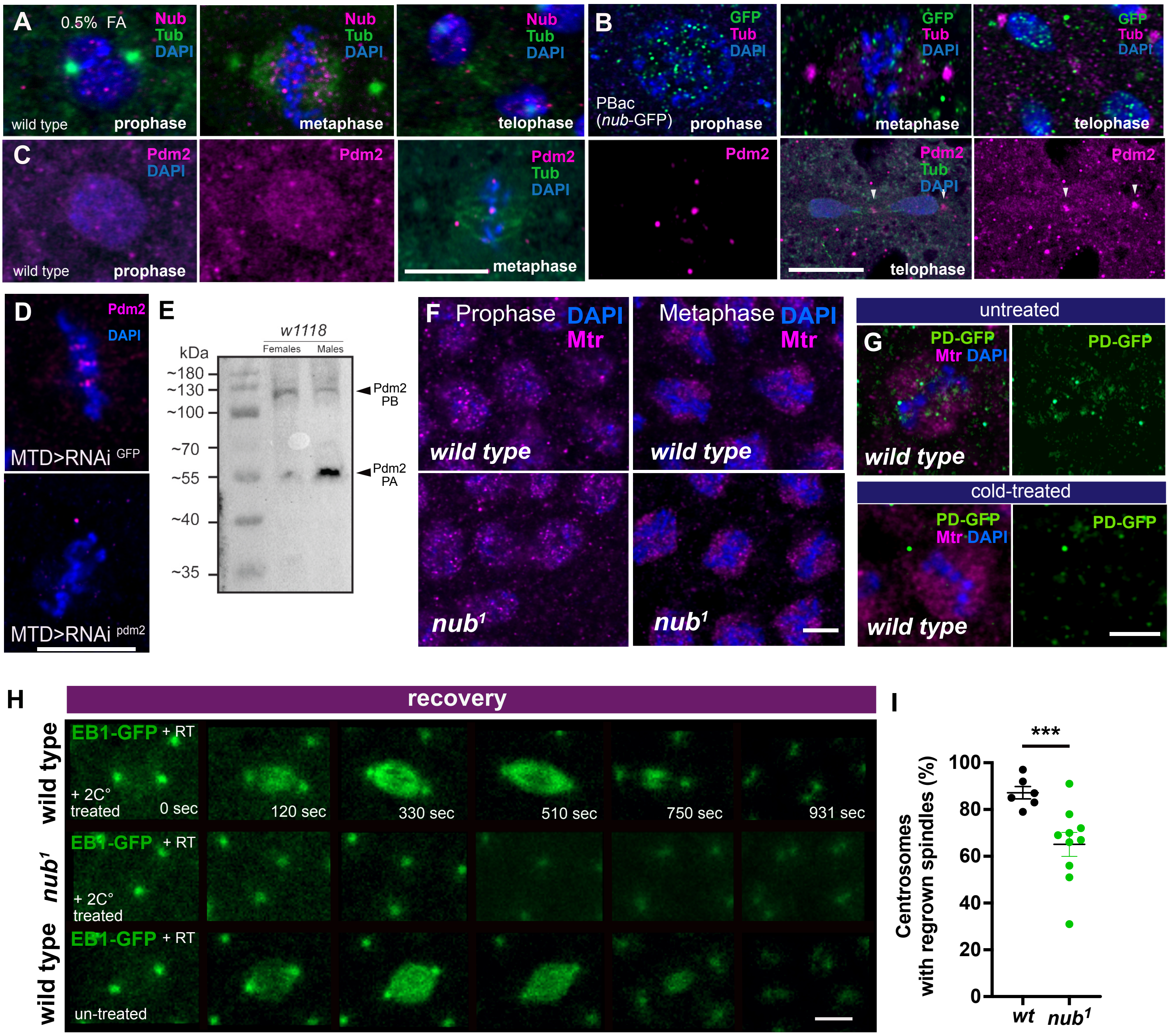

**Figure S6. Loss of Nub-PD affects spindle-MT regrowth but not spindle matrix structure.**

(A) Airy-scan confocal images of wild-type syncytial embryos stained with anti-Nub (magenta), anti- α- and γ-tubulin (green) and DNA with DAPI (blue) during prophase, metaphase and telophase. Embryos were fixed in 0.5% formaldehyde.
(B) Airy-scan confocal images showing the prophase, metaphase and telophase of a dividing nuclei of PBac (nub-GFP) syncytial embryo stained for anti-GFP (green) and DNA with DAPI (blue).
(C) Airy-scan confocal images showing the prophase, metaphase and telophase of a dividing nuclei of a wild type syncytial embryo stained with anti-Pdm2 (magenta), anti- β- & γ- tubulin (green) and DAPI (blue) for detecting DNA. Arrowheads indicate the mid-zone formed between two sister nuclei and the centrosomes.
(D) Confocal images of dividing nuclei in control MTD>RNAiGFP and MTD>RNAi pdm2 early embryos stained for Pdm2 (magenta) and DAPI (blue). Pdm2 signal is strongly reduced upon pdm2 RNAi indicating the specificity of the antibody.
(E) Western blot of fly extracts from w1118 adults showing the two endogenous Pdm2 isoforms (PA and PB). Both isoforms are present in female and male extracts.
(F) Airy-scan confocal images showing the prophase and metaphase of dividing nuclei of wild type and nub1 syncytial embryos stained for anti-Megator (Mtr, magenta) and DAPI (blue).
(G) Confocal images showing the metaphase of sfGFP-NubPD spindles in cold-treated or untreated embryos stained for anti-Megator (Mtr, magenta), GFP (green), and DAPI (blue).
(H) Airy-scan confocal images from time-lapse recordings showing the spindle-MT regrowth at room temperature, of cold treated (+2°C) or un-treated, live wild type and nub1 syncytial embryos expressing EB1-GFP (green).
(I) Plot showing the average pair of centrosomes without spindles (% per embryo) in cold-treated wild type (n = 6) and nub1 (n = 10 embryos). **** $p < 0.0001$. Statistical significance was calculated by t -test. Scale bars, 10μm.

Figure S6

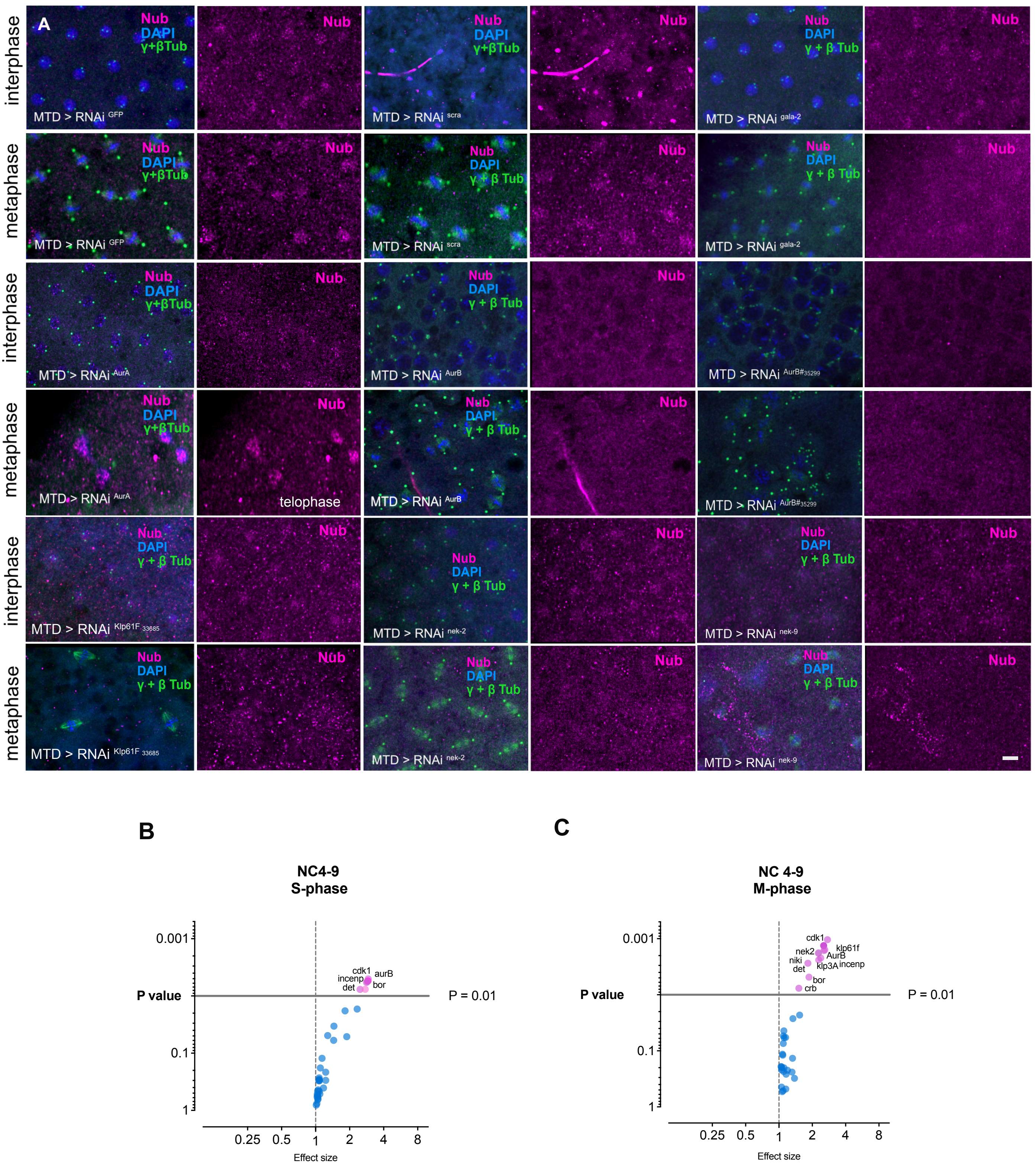
A
interphase
metaphase
interphase
metaphase
interphase
metaphase
Nub
DAPI
γ+βTub
MTD > RNAi GFP
MTD > RNAi scra
MTD > RNAi gala-2
MTD > RNAi AurA
MTD > RNAi AurB
MTD > RNAi AurB#35299
telophase
MTD > RNAi Klp61F 33685
MTD > RNAi nek-2
MTD > RNAi nek-9
B
NC4-9
S-phase
P value
0.001
0.1
1
P = 0.01
cdk1
aurB
incenp
det
bor
0.25
0.5
1
2
4
8
Effect size
C
NC 4-9
M-phase
cdk1
klp61f
nek2
AurB
niki
det
klp3A
incenp
bor
crb

Figure S7

**Figure S7. Nub-PD mitotic localization upon RNAi-mediated depletion of components of the mitotic machinery**

(A) Confocal images showing the cortex in syncytial embryos of *MTD-GAL4>RNAi*$^{GFP}$ , *MTD-GAL4>RNAi*$^{scra}$ , *MTD-GAL4>RNAi*$^{gala\text{-}2}$ , *MTD-GAL4>RNAi*$^{AurA}$ , *MTD-GAL4>RNAi*$^{AurB}$, *MTD-GAL4>RNAi*$^{AurB\#35299}$, *MTD-GAL4>RNAi*$^{klp61F\#33685}$*,* *MTD-GAL4>RNAi*$^{nek\text{-}2}$, *MTD-GAL4>RNAi*$^{nek\text{-}9}$ immunostained for *α*- and *γ*-tubulin (green), Nub (magenta) and DNA with DAPI (blue). The mitotic phases of the dividing nuclei are shown (prophase, metaphase). Nuclei in *MTD>AurA RNAi* were only possible to be captured in prophase and anaphase, as indicated.

(B-C) Volcano plots of statistical significance versus effect size for relative fluorescence intensity (RFI) values of Nub-PD in interphase nuclei in S-phase (B) and around the mitotic spindle in M-phase (C) during NC4-9. Dots (magenta) above the black line correspond to RNAi lines with significantly lower RFI values ($P < 0.01$) compared to control (*GAL4>RNAi*$^{GFP}$), indicating reduced or mislocalized Nub-PD. Dots below the significant threshold denote no difference compared to wild type. Scale bars, 10 µm.

### Supplementary Videos Legends

#### Videos S1-2. Fluorescence live imaging of syncytial nuclear divisions

Time-lapse recordings of a wild type (Video 1) and *nub*[1] (Video 2) embryos at NC11-13. Embryo are expressing the His2A-RFP reporter. Time indicates minutes (min). Videos are related to Fig. 3.

#### Video S3. Confocal live imaging of nuclear divisions in early embryos

Airy scan confocal time-lapse recordings of a wild type (left), *nub*[1] (right) embryos at NC11-13, expressing the His2A-RFP reporter. Time indicates seconds (sec). Several nuclei (indicted in magenta color) exhibit chromosome segregation defects and will eventually fall into the interior of the embryo for degradation, related to Fig. 3.

#### Video S4. Fluorescence live imaging of syncytial nuclear divisions

Time-lapse recordings of an anti-Nub injected wild type embryo at NC11-13. Embryo is expressing the nuclear reporter His2A-RFP. Time indicates minutes (min). The video is related to Fig. 3 and S3.

#### Videos S5-6. Confocal live imaging of spindles during nuclear divisions

Airy scan confocal time-lapse recordings of wild type (Video S5) and *nub*[1] (Video S6) embryos at NC11-13. Embryos express the EB1-GFP reporter, which traces microtubule plus-ends. *nub*[1] embryos exhibited disorganized spindles, with detached microtubule bundles often fusing with neighboring spindles or free centrosomes (Video S6), leading to clusters of multipolar spindles, related to Fig. 4.

**Supplementary Table S1.** Summary of mutations analyzed for Nub localization during interphase and metaphase.

| ***FlyBase ID*** | ***Gene*** | ***Mammal. homolog*** | ***Function*** | ***RNAi or mutant lines*** | ***RFI*** | | | |
|---|---|---|---|---|---|---|---|---|
| | | | | | **NC 4-9** | | **NC 10-12** | |
| | | | | | **S** | **M** | **S** | **M** |
| **NA** | *GFP* | | control | #41551 | 3.06 ± 0.14 | 2.16 ± 0.28 | 2.34 ± 0.35 | 1.96 ± 0.36 |
| **NA** | *$w^{1118}$* | | control | #3605 | 2. 91 ± 0.35 | 2.06 ± 0.31 | 2.15 ± 0.57 | 1.87 ± 0.22 |
| **FBgn0000147** | *AurA* | AurA | serine/threonine protein kinase | #41600 | 2.99 ± 0.58 | NA | 2.01 ± 0.68 | 1.76 ± 0.61 |
| **FBgn0000147** | *AurA* | AurA | serine/threonine protein kinase | #35763 | 2.86 ± 0.44 | 2.1 ± 0.34 | 1.92 ± 0.25 | 1.76 ± 0.11 |
| **FBgn0000063** | *Mps1* | Ttk | protein kinase | #35283 | 2.89 ± 0.22 | 2.12 ± 0.13 | 2.04 ± 0.24 | 1.87 ± 0.38 |
| **FBgn0000063** | *Mps1* | Ttk | protein kinase | #36658 | NA | NA | NA | NA[1] |
| **FBgn0033845** | *Mars* | Dlgap5 | microtubule-associated protein | #33929 | 2.94 [2] ± 0.65 | NA [1] | NA [1] | NA [1] |
| **FBgn0004378** | *klp61F III* | Kif11 | Kinesin-like protein at 61F | #33685 | 2.89 ± 0.28 | **1.48*** ± 0.09** | 2.26 ± 0.48 | **1.32*** ± 0.06** |
| **FBgn0004378** | *Klp61F II* | Kif11 | Kinesin-like protein at 61F | #35804 | 2.06 ± 0.35 | **1.39*** ± 0.02** | 1.87 ± 0.47 | **1.57*** ± 0.11** |
| **FBgn0004380** | *klp64D* | Kif3A | Kinesin-like protein at 64D | #40945 | | | | |
| **FBgn0011606** | *klp3A* | Kif4 | Kinesin-like protein at 3A | #40944 | 2.90 ± 0.19 | **1.52*** ± 0.13** | 2.47 ± 0.28 | **1.78** ± 0.21** |
| **FBgn0011606** | *klp3A* | kif4 | Kinesin-like protein at 3A | #43230 | 3.12 ± 0.17 | **1.77*** ± 0.13** | 2.29 ± 0.21 | **1.98** ± 0.31** |
| **FBgn0030268** | *klp10A* | Ttk | Kinesin-like protein at 10A | #33963 | 2.78 ± 0.38 | NA[1] | NA[1] | NA[1] |
| **FBgn0034824** | *klp59C* | Kif2C | Kinesin-like protein at 59C | #35596 | 2.61 ± 0.15 | 1.93 ± 0.42 | 2.14 ± 0.31 | 1.82 ± 0.20 |
| **FBgn0002924** | *ncd* | Kifc1/Kifc5b | kinesin-related microtubule motor protein | #58144 | 3.21 ± 0.05 | 2.16 ± 0.42 | 2.30 ± 0.24 | 1.81 ± 0.22 |
| **FBgn0261797** | *Dynein heavy chain 64C* | Dync1h1 | heavy chain subunit of the dynein motor complex | #36583 | 2.72 ± 0.34 | 1.69 ± 0.22 | 2.1 ± 0.38 | 1.89[1] ± 0.19 |
| **FBgn0261797** | *Dynein heavy chain 64C* | Dync1h1 | heavy chain subunit of the dynein motor complex | #36698 | 3.01 ± 0.29 | 1.98 ± 0.37 | 2.02 ± 0.37 | 1.85 ± 0.45 |

| | | | | | | | | |
|---|---|---|---|---|---|---|---|---|
| **FBgn0261797** | *Dynein-hc-64C(6-10]* | Dync1h1 | heavy chain subunit of the dynein motor complex | #8747 | 2.89 ± 0.15 | NA | 2.56 ± 0.40 | 1.99 ± 0.7 |
| **FBgn0261797** | *Dynein-hc-64C(6-6]* | Dync1h1 | heavy chain subunit of the dynein motor complex | #32015 | 2.87 ± 0.26 | 1.84 ± 0.32 | 2.01[1] ± 0.67 | NA[2] |
| **FBgn0261797** | *Dynein-hc-64C(6-6)/hc-64C6-10]* | Dync1h1 | heavy chain subunit of the dynein motor complex | #32015/+; #8747/+ | NA | 1.78 ± 0.44 | 1.89 ± 0.81 | 1.92[1] ± 0.68 |
| **FBgn0260991** | *Incenp* | INCENP | CPC member | #35366 | **1.29*** ± 0.16** | **1.16*** ± 0.38** | **1.38*** ± 0.21** | **1.19*** ± 0.18** |
| **FBgn0024227** | *AurB* | AurKC | serine-threonine kinase/ CPC member | #35299 | **1.19*** ± 0.11[1]** | **1.08*** ± 0.05[1]** | **1.19*** ± 0.11 [1]** | **NA [1]** |
| **FBgn0024227** | *AurB* | AurKC | serine-threonine kinase/ CPC member | #58308 | **1.15** ± 0.12** | **1.02*** ± 0.09** | **1.12** ± 0.10** | **1.24** [2] ± 0.14** |
| **FBgn0032105** | *Borealin-L* | borealin-related | DNA binding protein/CPC member | #56942 | **2.65* ± 0.27** | **1.62*** ± 0.19** | **1.52* ± 0.34** | **1.64** ± 0.21** |
| **FBgn0264291** | *Deterin* | Survivin/ BIRC5) | CPC member | #36612 | 2.73[1] ± 0.57 | NA [1] | NA [1] | NA [1] |
| **FBgn0264291** | *Deterin* | Survivin/ BIRC5) | CPC member | #36600 | **1.49** ± 0.19** | **1.28*** ± 0.04** | **1.12*** ± 0.11** | NA [1] |
| **FBgn0029970** | *nek2* | Nek2 | protein kinase | #35328 | 2.71 ± 0.32 | **1.55 *** ± 0.16** | 2.35 ± 0.46 | **1.88** ± 0.46** |
| **FBgn0029970** | *nek2* | Nek2 | protein kinase | #60106 | 2.26 ± 0.11 | **2.13 ** ± 0.62** | **2.79 * ± 0.43** | **2.61* ± 0.57** |
| **FBgn0045980** | *NiKi* | Nek9 | serine/threonine kinase | #53735 | 2.54 ± 0.49 | **1.36*** ± 0.11** | 2.68 ± 0.39 | **1.14*** ± 0.04** |
| **FBgn0036107** | *Galla-2* | CIAO2B | Cytoplasmic/CGX complex member | #58320 | 2.87 ± 0.52 | 1.80 ± 0.4 | 2.19 ± 0.71 | 2.41 ± 0.68 |
| **FBgn0259685** | *Crb* | Crb1-3 | Transmembrane protein | #38903 | 2.75 **± 0.72** | **1.72** ± 0.32** | 1.95 **± 0.62** | **1.39** ± 0.11** |
| **FBgn0261850** | *Xpd* | Ercc2 | DNA helicase /CGX complex member | #65883 | 2.68 **± 0.81** | 1.70 **± 0.57** | 1.91 **± 0.84** | 1.88 **± 0.61** |
| **FBgn0000826** | *png* | Nek2 | Ser/Thr kinase/ mRNA translation | #55861 | NA | NA | NA | NA[2] |
| **FBgn0016070** | *smg* | Samd4 | sequence-specific RNA-binding protein | #35477 | NA | 1.53 ± 0.49 | **1.61* ± 0.37** | NA [2] |
| **FBgn0261385** | *scra /Anillin* | Anln | actin binding. protein | #41841 | 2.82 ± 0.72 | 1.72 ± 0.43 | 1.89 ± 0.39 | 1.84 |
| **FBgn0046706** | *Haspin* | Haspin | Serine/threonine-protein kinase | #35276 | 2.76 ± 0.53 | 1.77 ± 0.56 | 1.98 ± 0.33 | 1.87 ± 0.49 |

| | | | | | | | | |
|---|---|---|---|---|---|---|---|---|
| **FBgn0003890** | *βTub97EF* | Tubb2a | *β*-tubulin protein | #64858 | 2.98 ± 0.68 | 2.21 ± 0.56 | 1.98 ± 0.62 | 1.87 ± 0.44 |
| **FBgn0261278** | *grp/Chk1* | CHEK1 | serine/threonine protein kinase | #36685 | NA[2] | NA[2] | NA[2] | NA[2] |
| **FBgn0261278** | *grp/Chk1* | CHEK1 | serine/threonine protein kinase | #62155 | 2.90 ± 0.88 | 2.04 ± 0.56 | 1.67 ± 0.31 | 1.72 ± 0.42 |
| **FBgn0266282** | *SMC6$^{Del}$* | SMC6 | subunit of the SMC5/6 complex | NA (Tran *et al*, 2016) | NA | 2.91 ± 0.66 | 1.82 ± 0.32 | NA[1] |
| **FBgn0004106** | *Cdk12* | *Cdk13* | cyclin-dependent kinase | #42775 | | | | |
| **FBgn0004106** | *Cdk1* | *Cdk1* | cyclin-dependent kinase | #35350 | **1.19*** ± 0.10** | NA[1] | NA[1] | **1.79*** ± 0.31** |
| **FBgn0004106** | *Cdk1* | *Cdk1* | cyclin-dependent kinase | #65396 (WT) [3] | **1.05*** ± 0.08** | **1.27*** ± 0.10** | **1.28*** ± 0.16** | **1.51** ± 0.23** |
| **FBgn0004106** | *Cdk1* | *Cdk1* | cyclin-dependent kinase | #65394 [3] (Active) | **1.12 ± 0.24** | **1.41 ± 0.18** | **1.26 ± 0.07** | **1.32 ± 0.14** |
| **FBgn0004106** | *Cdk1* | *Cdk1* | cyclin-dependent kinase | #65398 [3] (Active) | **1.46 ± 0.11** | **1.39 ± 0.17** | **1.57 ± 0.33** | **1.12 ± 0.08** |
| **FBgn0004106** | *Cdk1* | *Cdk1* | cyclin-dependent kinase | #65395 (Active) [3] | NA[2] | NA[2] | NA[2] | NA[2] |

| p>0.05 | p<0.05 | p<0.01 | p<0.0001 |
|---|---|---|---|

Each gene analyzed is presented with its synonym, homolog in mammals and function according to FlyBase. The unique Flybase identification number (ID) is shown on the first column. The stock number or source of RNAi lines or mutants are indicated. *RFI*: relative fluorescence intensity (RFI) of the Nub is shown. *RFI= fluorescence intensity of Nub in nucleus or spindles/cytoplasm*. Values are means ± SD. RFI values significantly different than the control or wild type are highlighted in bold. NA: not available; NC: nuclear cycle; S: DNA synthesis phase; M: Mitotic phase; (n > 5 embryos in every condition).

[1] Early-embryonic lethality with incomplete penetrance
[2] Early-embryonic lethal
[3] UASp over-expression
*, $p < 0.05$, **, $p < 0.01$ ***, $p < 0.0001$

**Supplementary Table 2.** List of key resources used in the study.

| **Reagent or resource** | **Source or reference** | **Identifiers** |
|---|---|---|
| **Experimental Models: Organisms/strains** | | |
| *$w^{1118}$* | Bloomington Drosophila Stock Center (BDSC), Indiana University | RRID:BDSC_5905 |
| *$nub^{1}$* mutant | BDSC | RRID:BDSC_357 |
| His2Av-RFP | BDSC | RRID:BDSC_23650 |
| EB1-GFP | BDSC | RRID:BDSC_57327 |
| Klp61F-GFP | BDSC | RRID:BDSC_35509 |
| PBac{nub-GFP) | BDSC | RRID:BDSC 96785 |
| mat-gal4 | BDSC | RRID:BDSC_7062 |
| mat-gal4 | BDSC | RRID:BDSC_7063 |
| MTD-gal4 | BDSC | RRID:BDSC_31777 |
| ***sfGFP-NubPB*** | This study | NA |
| ***sfGFP-NubPD*** | This study | NA |
| *nos-deGradFP* | Gift from Melissa Harrison University of Wisconsin-Madison | NA |
| *hb*-deGradFP | BDSC | RRID:BDSC_96802 |
| CH321-40A01 (*nub* duplication) | BDSC | RRID:BDSC_89924 |
| *$tud^{1}$* | BDSC | RRID:BDSC_1786 |
| *nub-RNAi* | BDSC | RRID:BDSC_55305 |
| GFP-RNAi | BDSC | RRID:BDSC_41551 |
| **Experimental Models: Cell lines** | | |
| *D. melanogaster* S2 cell line | | RRID:CVCL_Z232 |
| *D. melanogaster* S2 cell-stably expressing Histone2B-GFP and tubulin-mCherry | Afonso et al. 2014. [1] | NA |
| Human cell line: HeLa cell | Sigma-Aldrich, | Cat# ECACC, 93021013 |
| Human cell line: HEK293 | ATCC | Cat# CRL-1573 |
| **Primary antibodies** | | |

| | | |
|---|---|---|
| Rabbit anti-Nub/Pdm1 | Gift from Xiaohang Yang and Cai Yu, Temasek Life Sciences Laboratory, National University of Singapore, Singapore | |
| Mouse monoclonal anti-Nub | Development studies hybridoma bank (DSHB) | Cat# Nub 2D4, RRID:AB_2722119 |
| Rabbit Polyclonal anti-Nub (peptide antibody) | Dantoft et al. 2013. [3] | NA |
| Mouse Monoclonal anti-β-tubulin | DSHB | Cat# E7, RRID:AB_528499 |
| Mouse Monoclonal anti-γ-tubulin | Sigma-Aldrich | Cat# T5326, RRID:AB_532292 |
| Rabbit Monoclonal anti-GFP | Thermo Fisher Scientific | Cat# A-11122 RRID:AB_221569 |
| Rabbit Polyclonal anti-Pdm2 (peptide antibody) | This study | NA |
| Rabbit Polyclonal anti-PH3 | Millipore | Cat# 06-570, RRID:AB_310177 |
| Mouse anti-Megator | DSHB | Cat# 12F10-5F11, RRID:AB_2721935 |
| Mouse monoclonal anti-Lamin-B | DSHB | ADL84 RRID:AB_528338 |
| **Secondary antibodies** | | |
| Goat anti-rabbit Alexa fluor 488 | Thermo Fisher Scientific | Cat# A66785, RRID:AB_3251385) |
| Goat anti-mouse Alexa fluor 647 | Thermo Fisher Scientific | Cat# A-21235, RRID:AB_2535804 |
| Goat anti-rabbit Alexa fluor 594 | Thermo Fisher Scientific | Cat# A-11012, RRID:AB_2534079 |
| Donkey anti-Mouse Alexa fluor 488 | Thermo Fisher Scientific | Cat# A-21202, RRID:AB_141607 |
| Cy3 | Jackson Immunochemicals | Variable host and target species |
| Cy5 | Jackson Immunochemicals | Variable host and target species |

| | | |
|---|---|---|
| **Chemicals** | | |
| Heptane | Sigma | Cat. #H2198 |
| Sodium Hypochlorite | VWR | Cat. #27900.296 |
| TRIzol reagent | Thermo fisher scientific | Cat. #15596018 |
| Schneider's media | Thermo Fisher scientific/Gibco | Cat. #21720024 |
| DMEM F12 | Thermo Fisher scientific/Gibco | Cat. #12634010 |
| Penicillin-streptomycin antibiotics | Thermo Fisher scientific/Gibco | Cat. #15140122 |
| DMEM GlutaMAX | Thermo Fisher scientific/Gibco | Cat. #31966021 |
| Halocarbon oil (#700) | Sigma | Cat. # H8898 |
| ViaFluor 488 | Biotium | Cat. #70062 |
| Fetal Bovine Serum | Thermo Fisher/Gibco | Cat. #A5670801 |
| NGS | Thermo Fisher Scientific | Cat. #31873 |
| Formaldehyde 37% | | Cat. # 47608 or F8775 |
| VECTASHIELD PLUS Antifade | BioNordika | Cat. #VEH-1900-10 |
| **Commercial assay kits** | | |
| Mini-prep kit | QIAGEN | Cat. #27106 |
| Effectene transfection kit | QIAGEN | Cat. #301425 |
| Lipofectamine™ RNAiMAX transfection reagent | Thermo Fisher Scientific | Cat. #13778100 |
| LR clonase kit | Thermo Fisher Scientific | Cat. #11791020 |
| SuperScript IV First-strand Synthesis System | Thermo Fisher Scientific | Cat. #18091050 |
| T7 RiboMAX™ Express Large Scale RNA Production System | Promega | Cat. #P1320 |
| pENTR™/D-TOPO™ Cloning Kit | Thermo Fisher Scientific | Cat. #K240020SP |
| Real-time PCR assay to detect Mycoplasma spp. | IDEXX BioAnalytics | |
| **Primers** | | |
| Nub-PD.HL.Fw | atgcac***ctcgag***cacacacccatacactgacaac | |

| | | |
|---|---|---|
| Nub-PD.HL.Rw | tggtac***aagctt***TAGCTCCGACATAACCATTTTC | |
| Nub-PDsfGFP.fw | tGTGAAAATGGTTATGTCGGAGCTAGTGTCCAAGGGCGAGGAG | |
| Nub-PDsfGFP.Rw | ccTCGGGACTAGCGGTGTGCCAACGGATTATCTTTCTAGGGTTAATCTTG | |
| Nub-PD.HR.Fw | atgcac***aagctt***CGTTGGCACACCGCTAGTCC | |
| Nub-PD.HR.Rw | tggtac***GGATCC***gttctcgcctggcaccgcca | |
| Nub-PB.HL.Fw | atgcac***ctcgag***ggcttgcgatcatgaatattcag | |
| Nub-PB.HL.Rw | tggtac***aagctt***GGTGGCCTCGCCATTGATTTTGTATTTC | |
| sfGFP.RBfw | ATACAAAATCAATGGCGAGGCCACCGTGTCCAAGGGCGAGGAG | |
| sfGFP.RBrw | GCAGTGTGGATGCTGAAACGCCTTCGATTATCTTTCTAGGGTTAATCTTG | |
| Nub-PB.HR.Fw | atgcac***ggatcc***GAAGGCGTTTCAGCATCCAC | |
| Nub-PB.HR.Rw | tggtac***GCGGCCGC***gacacacacgttgccaaaac | |
| Nub-RB.Trg1_sense | **GTCG**GGATGCTGAAACGCCTTCGG | |
| Nub-RB.Trg1_antisense | **AAAC**CCGAAGGCGTTTCAGCATCC | |
| Nub-RB.Trg2_sense | **GTCG**CGGATGATGATACAGCAGTG | |
| Nub-RB.Trg2_antisense | **AAAC**CACTGCTGTATCATCATCCG | |
| Nub-RD.Trg1_sense | **GTCG**TGGTTATGTCGGAGCTACGT | |
| Nub-RD.Trg1_antisense | **AAAC**ACGTAGCTCCGACATAACCA | |
| Nub-RD.Trg2_sense | **GTCG**TTGGCACACCGCTAGTCCCG | |
| Nub-RD.Trg2_antisense | **AAAC**CGGGACTAGCGGTGTGCCAA | |
| **Plasmids/DNA** | | |
| pCFD3-dU6:3gRNA | Addgene | RRID:Addgene_49410 |

| | | |
|---|---|---|
| pBS II SK(+) | Addgene | Cat. #212205 |
| pScarlessHD-sfGFP-DsRed | Addgene | RRID:Addgene_80811 |
| **Oligonucleotides** | | |
| Human POU2F1/Oct1 siRNA | Santa Cruz Biotechnology | Cat. #sc-36119 |
| Control siRNA | Santa Cruz Biotechnology | Cat. #sc-37007 |
| **Software** | | |
| ImageJ | National Institute of Health, USA https://imagej.nih.gov/ij/ | RRID:SCR_002285 |
| GraphPad Prism 10.1 | GraphPad Prism https://www.graphpad.com/scientific-software/prism/ | RRID:SCR_002798 |
| ZEN Blue or Zen2011 | ZEISS ZEN Microscopy Software https://www.zeiss.com/corporate/int/home.html | RRID:SCR_013672 |
| Affinity designer | Serif (Europe) Ltd https://affinity.store/en-us/ | RRID:SCR_016952 |
| JMP Pro 17.0 statistical program | SAS Institute | RRID:SCR_022199 |
